# Identification of Potent Small Molecule Inhibitors of SARS-CoV-2 Entry

**DOI:** 10.1101/2021.08.05.455262

**Authors:** Sonia Mediouni, Huihui Mou, Yuka Otsuka, Joseph Anthony Jablonski, Robert Scott Adcock, Lalit Batra, Dong-Hoon Chung, Christopher Rood, Ian Mitchelle S. de Vera, Ronald Rahaim, Sultan Ullah, Xuerong Yu, Tu-Trinh Nguyen, Mitchell Hull, Emily Chen, Thomas D. Bannister, Pierre Baillargeon, Louis Scampavia, Michael Farzan, Susana T. Valente, Timothy P. Spicer

**Affiliations:** Scripps Research, Department of Immunology and Microbiology, Scripps Research, Jupiter, FL 33458, USA; Scripps Research, Department of Molecular Medicine, Scripps Research, Jupiter, FL 33458, USA; Center for Predictive Medicine, Department of Microbiology Immunology, School of Medicine, University of Louisville, KY 40202, USA; Department of Pharmacology and Physiology, Saint Louis University School of Medicine, St. Louis, MO 63104, USA; CALIBR, Scripps Research, 11119 N Torrey Pines Rd, La Jolla, CA 9203, USA

## Abstract

The severe acute respiratory syndrome coronavirus 2 responsible for COVID-19 remains a persistent threat to mankind, especially for the immunocompromised and elderly for which the vaccine may have limited effectiveness. Entry of SARS-CoV-2 requires a high affinity interaction of the viral spike protein with the cellular receptor angiotensin-converting enzyme 2. Novel mutations on the spike protein correlate with the high transmissibility of new variants of SARS-CoV-2, highlighting the need for small molecule inhibitors of virus entry into target cells. We report the identification of such inhibitors through a robust high-throughput screen testing 15,000 small molecules from unique libraries. Several leads were validated in a suite of mechanistic assays, including whole cell SARS-CoV-2 infectivity assays. The main lead compound, Calpeptin, was further characterized using SARS-CoV-1 and the novel SARS-CoV-2 variant entry assays, SARS-CoV-2 protease assays and molecular docking. This study reveals Calpeptin as a potent and specific inhibitor of SARS-CoV-2 and some variants.

## INTRODUCTION

The severe acute respiratory syndrome coronavirus 2 (SARS□CoV□2) is the pathogen responsible for the COVID-19 disease which reached pandemic designation in early 2020. Even with the release of FDA-approved vaccines in early 2021, the pandemic remains the number one worldwide public health threat with massive negative social and economic impacts. Much of the world still faces widespread spikes in confirmed cases, hospitalizations, demands on critical care, and mortality^1^. The highly infectious delta variant has become the predominant strain in the United States. Those that remain vaccine resistant are taking the brunt of infections but, breakthrough cases are occurring more frequently. Infection rates vary by location, but in almost all areas, episodes of disease reemergence have been observed. People with underlying medical conditions, especially elderly that are immunosuppressed, have been heavily impacted. In fact, adults over 65 years of age represent 80% of hospitalizations with a 23-fold higher risk of death^2^.

Common symptoms include high fever, cough, fatigue, breathing difficulties, and loss of taste and smell. Severe cases also exhibit clinical manifestations such as pneumonia, acute respiratory distress syndrome and clotting disorders^3, 4^.The disease spread is primarily *via* micro droplet dispersion, typically by inhalation, and thus measures that reduce or eliminate exposures to infected people are highly recommended. These include regular wearing of masks, social distancing, avoiding groups of people other than the close family unit, self-isolation, and avoidance of indoor spaces with substandard ventilation^5–7^. Such preventive measures, in some cases, have been aided by electronic apps to identify exposures^5^, which seem effective yet compliance over prolonged periods tends to wane prompting additional waves of COVID-19 and thus reinvigorating the overburdening of the health care system, economic regression, and social instability.

Massive worldwide efforts towards the identification of novel treatments against SARS-CoV-2 are underway. Development of safe drugs with novel small molecules is laborious and time consuming; hence, many groups including our own have been investigating the repurposing of well-established drugs. For instance, the antiviral drug Remdesivir, a nucleotide analog with a broad-spectrum antiviral activity was used early into the pandemic in several countries despite its modest benefits and the burden of its intravenous administration^8^. Similarly, the anti-malarial drug chloroquine or its derivative, hydroxychloroquine, was tested and had mixed results and multiple safety concerns prompting its emergency use authorization to be revoked by the FDA^9^. Fortunately, at least three vaccines have been approved by the FDA, also for emergency use, which are currently being distributed broadly bringing hope to an end of the pandemic. However, with respect to vaccination, significant hurdles remain. Length of protection conferred upon vaccination isn’t fully understood, and in some cases when multiple doses are required, the spacing between vaccine doses that will confer the best immune response is still not clearly established. In addition, vaccine hesitancy is still a major concern, which is aggravated by the scarcity of vaccine doses in certain locations across the world. The appearance of novel highly infectious variants with mutations in the spike protein may also diminish the efficacy of the vaccine^10, 11^, reinforcing the need for additional improved therapeutics.

Two SARS-CoV-2 entry pathways have been described and are well understood. This is in part due to the close similarity of SARS-CoV-2 to the SARS-CoV-1, the causative agent for a global outbreak in 2002-2003^12^. Both entry pathways, being cell surface and endosomal, require the sequential function of two subunits, S1 and S2 of the spike protein (S), localized on the viral lipid membrane. The S1 subunit, *via* its receptor binding domain (RBD), binds a functional angiotensin-converting enzyme 2 (ACE2) receptor on the target cells^13, 14^. Next, the S2 subunit facilitates the fusion of the cell and viral membranes, *via* conformational changes followed by proteolytic activation at the S1/S2 boundary and dissociation of S1. The cleavage is performed either by the cell surface serine protease, transmembrane serine protease 2 (TMPRSS2) or the lysosomal protease, Cathepsin L (**Fig.1A**), according to the pathway taken. It is, however, unclear whether SARS-CoV-2 uses an additional co-receptor, but evidence is mounting in that direction^15^. In sum, inhibitors that block either the interaction of the viral S1 RBD with cellular receptors or the fusion mediated by S2 domain should inhibit entry of the virus.

**Fig.1.**
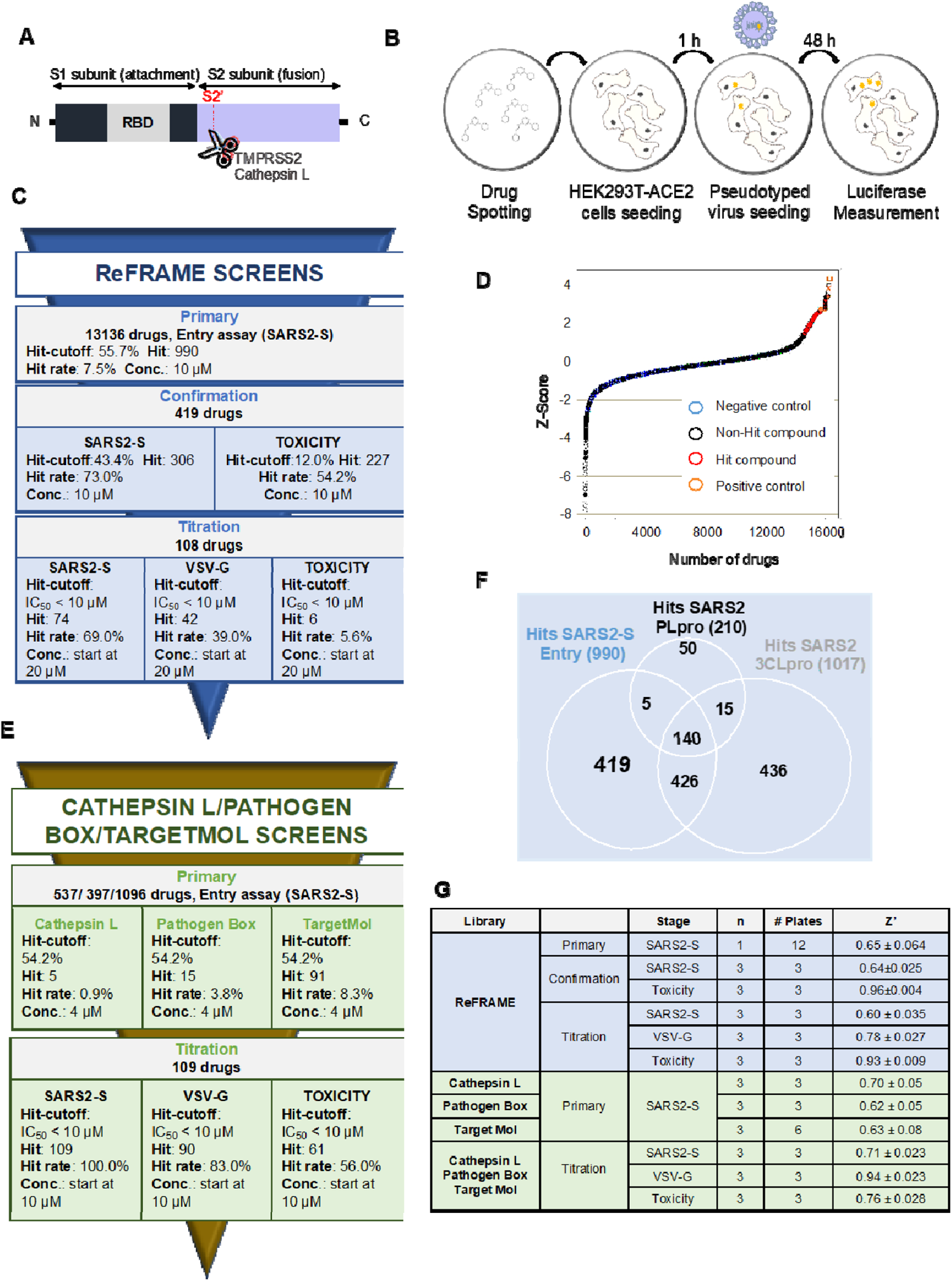
High-throughput of ReFRAME, Pathogen Box, TargetMol and Cathepsin L drug libraries for SARS-CoV-2 antiviral compounds. **A. Schematic of the spike protein of SARS-CoV-2**. RBD: receptor binding domain. **B. Schematic of the High-throughput assay**. Compounds were pre-spotted in 1536-well plates. Next, 2,000 HEK293T-ACE2 cells were added to each well and pre-incubated with each compound for 1 h, followed by infection with MLV reporter luciferase virus pseudotyped with the SARS-CoV-2 Spike protein (SARS2-S) or VSV-G protein (VSV-G). Luciferase was measured 48 hour later. **C. Summary of the ReFRAME library results.** Conc.: concentration. **D. Distribution of Z-Score for primary screens of each library.** Scatter plot of Z_Score for all samples tested from the ReFrame library (N=1; circle) and other libraries (N=3; Cathepsin L: square; Pathogen Box: cross; TargetMol: filled circle). Total of 16320 samples. Positive controls: orange; Negative control: cyan; Hit compounds: red; non-hit compounds: black. **E. Summary of the 3 other libraries results. F. ReFrame library screening against different targets: SARS2-S, 3CLpro and PLpro.** Venn diagram analysis of comparison between hits from SARS2-S entry, 3CLpro and PLpro assay against ReFRAME library results. There are 419 compounds that are SARS2-entry specific potential inhibitors**. G. Robustness in terms of Z’ score of each screen for each library.**

Based on this hypothesis, we undertook a High Throughput Screening (HTS) campaign using a comprehensive library of drugs that may be repurposed to seek potential therapeutics against SARS-CoV-2 infection, focusing specifically on the disruption of the entry process. We screened the ReFRAME library of 13,136 small molecules that have reached clinical development or have undergone preclinical profiling^16^, which increases our chance of identifying leads with well-established safety profiles and rapid progression to clinic. We also screened a collection of ~1,500 molecules of the Cathepsin L and TargetMol libraries, selected for their computational docking scores for interacting at the ACE2-RBD interface and drug-likeness. Furthermore, we screened a collection termed the Pathogen Box, consisting of ~400 compounds with known or suspected antiviral activity ^17^. We utilized a tiered approach: first screening the libraries against a Moloney Murine Leukemia virus (MLV) reporter luciferase virus pseudotyped with the SARS-CoV-2 S protein (SARS2-S), second the hits were confirmed in this same system, and third two counterscreens were performed with one using the Vesicular stomatitis virus G (VSV-G) enveloped virus to assess specificity and the other to gauge cytotoxicity. A series of potent and unique hits blocking virus entry emerged and which were validated in multiple mechanistic based assays. The lead compound, Calpeptin, was found to be a potent and specific inhibitor of SARS-CoV-2 including highly virulent variants.

## MATERIALS AND METHODS

### Data reporting

The number of samples tested was completely unbiased and all compounds were formatted in a randomized manner using an acoustic dispenser for analysis in the primary HTS campaigns. For the ReFRAME library, all structural information was withheld until primary hits had been validated. All other downstream experiments were not randomized nor blinded as structures were known and hits were chosen using statistical parameters described below.

### Plasmids

The plasmids for expression of the S or ACE2 proteins were created with fragments synthesized by Integrated DNA Technologies (IDT, Coralville, IA, USA). Fragment ligation was performed using In-Fusion® HD Cloning Kit (Takara Bio USA) according to manufacturer’s instructions. The expression plasmids including variant mutations were made by site-directed mutagenesis.

### Cell line

HEK293T cells (human embryonic kidney; ATCC CRL-3216, VA, USA) were maintained in growth medium composed of Dulbecco’s Modified Eagle Medium (DMEM, Life Technologies) supplemented with 2 mM Glutamine (Life Technologies), 1% non-essential amino acids (Life Technologies), 100 U/mL penicillin, 100 µg/mL streptomycin (Life Technologies) and 10% FBS (Sigma-Aldrich, St. Louis, MI, USA) at 37°C in 5% CO_2_. The HEK293T cell line expressing human ACE2 was created by transduction with VSV-G protein-pseudotyped MLV containing pQCXIP-myc-ACE2-c9, as previously described^18^. Briefly, HEK293T cells were co-transfected with three plasmids, pMLV-gag-pol, pCAGGS-VSV-G and pQCXIP-myc-ACE2-c9 and the medium was replaced the next day. The supernatant containing the pseudotyped virus was harvested 72 hours post-transfection and clarified by passing through 0.45 μm filter. The parental HEK293T cells were transduced with this virus and the HEK293T-ACE2 cell line was selected and maintained with medium containing puromycin (Sigma). ACE2 expression was confirmed by immunofluorescence staining using mouse monoclonal antibody against c-Myc antibody 9E10 (Thermo Fisher) and Goat-anti-mouse FITC (Jackson ImmunoResearch Laboratories, Inc). The HEK293T-ACE2 stable cells were maintained in growth medium, which contains 10% FBS (Sigma), 1% Penicillin-streptomycin (Sigma) and 3 µg/ml puromycin (Sigma) in DMEM (Corning).

An HEK293T cell line expressing human ACE2 and TMPRSS2 (HEK293T-ACE2-TMPRSS2) and Vero CCL81 cells were kindly provided by Dr. Choe (Scripps Research Florida, USA). They were maintained in growth media composed of DMEM (Life Technologies) supplemented with 10% FBS, 1/100 L-Glutamine: Penicillin: Streptomycin (GeminiBio). HEK293T-ACE2-TMPRSS2 cells were kept under selection with 1.3 µg/mL of puromycin (Sigma).

Vero E6 cells (ATCC CRL-1586) were obtained from ATCC and cultured in DMEM (HyClone.) with 10% FBS. Vero E6 cells enriched in ACE2 were selected by FACS with antibody against human ACE2. Vero E6 cells were stained with Goat anti-Human Phycoerythrin-conjugated ACE2 Polyclonal Antibody (R&D Systems) for 30 min at 4 °C in dark. Cells were then washed with HBSS and suspended in Pre-Sort buffer (BD Biosciences). ACE2+ and ACE2- were gated based on the Goat IgG Isotype Control (R&D Systems) and sorted on BD FACSAria^TM^ III. The ACE2 high and low enriched cell populations were then cultured once and then used for the experiment.

### Pseudotyped virus production and titration

MLVs pseudotyped with various envelope proteins were generated as previously described^19^. Briefly, HEK293T cells were co-transfected with three plasmids, pMLV-gag-pol, pQC-Fluc and pCAGGS-SARS2-SFmut-cflag or pCAGGS-VSV-G using PEI 40K (Polysciences, Inc.), and the medium was refreshed 6 hours later. The supernatant containing the pseudotyped virus was harvested 72 hours post-transfection and clarified through a 0.45 µm filter. Clarified viral stocks were supplemented with 10 mM HEPES and stored at −80 °C for long-term storage. Pseudotyped viruses were titrated using TCID_50_ method. Briefly, HEK293T-ACE2 cells were seeded in 96-well plates at 50-60% confluency upon observation the following day. Then, 50 µL of media was removed from each well and replaced with 50 µL of the serially diluted pseudotyped virus or media. The plate was centrifuged at 4°C, 3000g for 30 mins (spinoculation) and incubated for 2 hours at 37°C and 5% CO_2_^20^. Virus containing media was then aspirated and 100 µL of fresh media with 1 µg/mL puromycin was added to the wells. The plate was further incubated for 48 hours at 37°C and 5% CO_2_. Then, 100 µL/well of OneGlo (Promega) was added and luminescent signal was read. The average and 3 standard deviations of the signal level for media only wells were calculated and used as a cutoff. TCID_50_ was calculated using Reed & Muench Calculator^21^.

### SARS2-S entry Assay for HTS

Compounds were pre-spotted into 1536 well plates (Greiner BioOne part 789173-F) at either 5 nL for 10 mM stocks of compounds at CALIBR for the ReFRAME library or 20 nL for either 1 mM or 2.5 mM stocks of compounds for the Cathepsin L, Pathogen Box or TargetMol library, at Scripps Molecular Screening Center^16, 22^. The HEK293T-ACE2 cells were seeded at 2,000 cells/well, using an Aurora FRD (Aurora Discovery), at 2.5 µL/well. Plates were incubated for 1 hour at 37°C and 5% CO_2_. Pseudotyped virus was then dispensed at 2.5 µL/well, at a multiplicity of infection (MOI) of 0.1 for primary and confirmation assays and 0.5 for titration assays. After the 48-hour incubation period at 37°C and 5% CO_2_, plates were removed from the incubator and allowed to equilibrate at room temperature for 15 mins. OneGlo (Promega) Luciferase reagent was then added at 5 µL/well and incubated for 10 mins at room temperature. The luminescence was subsequently measured using a ViewLux (PerkinElmer) for 5 secs. The process is summarized in **Fig.1B**. High control wells contained HEK293T-ACE2 cells + Media + vehicle (DMSO), and the low control and data wells had HEK293T-ACE2 cells + SARS2-S + test compound or vehicle.

### Counterscreen Assays for HTS

The first counterscreen assay followed the same protocol as the SARS2-S entry assay but instead uses VSV-G pseudotyped virus, to identify non-specific entry inhibitors. In addition, and in parallel, the cytotoxicity of selected compounds was also tested during the campaign using CellTiter-Glo (Promega). The controls for the VSV-G assay followed the exact logic of the primary assay while the toxicity assay incorporated no cells (high control) *vs* cells treated with vehicle only (low control) as controls.

### Screening Libraries

*ReFRAME Library* - CALIBR, a division of Scripps Research, has partnered with the Bill and Melinda Gates Foundation to form an integrated platform of drug candidates. At the time of this effort the ReFRAME contained 13,136 purchased or resynthesized FDA-approved/registered drugs (~40%), as well as investigational new drugs currently or previously in any phase of clinical development (~60%).

*Pathogen Box Library* - This library is comprised of 400 diverse, drug-like molecules active against neglected diseases of interest. It was provided by the Medicines for Malaria Venture.

*TargetMol Library* - TargetMol performed a CADD *in silico* docking study using the Swiss-Model Homology Modelling process to generate reliable protein models or 3D protein structures of the RBD of S protein and ACE2. Based on these protein structures they identified 462 top ranked for ACE2 by molecular docking virtual screening against 15,376 compound structures.

*Cathepsin L library* – As part of the Molecular Libraries Probe Center Network, we had previously screened the target cathepsin L1 and had identified 1482 active molecules (see PubChem AID 1906) from a total of 302,755 small molecules in the NIH repository. We then surveyed our Scripps Drug Discovery Library of greater than 665K small molecules to identify ~ 450 compounds that overlapped with a Tanimoto score >80% identity with these inhibitors. These compounds were cherry-picked and registered into source plate for screening.

### Screening data acquisition, normalization, representation, and analysis for HTS

Data files were uploaded into the Scripps institutional HTS database (Symyx Technologies, Santa Clara, CA) for plate QC and hit identification. Activity for each well was normalized on a per-plate basis using the following equation:

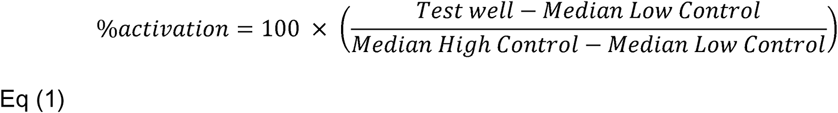

where “High Control” represents wells containing HEK293T-ACE2 cells + assay Media + vehicle (DMSO) wells, while “Low Control” represents wells containing HEK293T-ACE2 cells + SARS2-S + vehicle and finally the “Data Wells” contain HEK293T-ACE2 cells + SARS2-S + test compounds. The Z’ and S:B were calculated using the High Control and Low Control wells. A Z’ value greater than 0.5 was required for a plate to be considered acceptable^23^. Z-score was also calculated to show the distribution of the well-to-well results by using following equation:

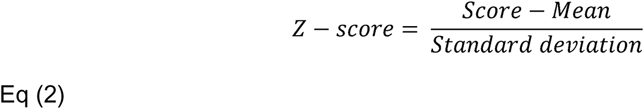

A positive Z-score indicates the raw score is higher than average^24^.

### Follow-up assays

HEK293T-ACE2 cells, HEK293T-ACE2-TMPRSS2 cells or Vero CCL81 cells were seeded at 1-1.5×10^4^ cells/well in a 96-well plate. The next day, compounds (or DMSO control) were added to cells at multiple concentrations, in presence of pseudotyped viruses. This was done separately using SARS2-S, different mutants of SARS2-S, VSV-G and finally with pseudotyped virus containing the S protein from SARS-CoV-1 “SARS1-S”. The MOI of each virus was chosen based on an equivalent level of luciferase production. After a 48-hour incubation at 37°C and 5% CO_2_, plates were removed from the incubator and allowed to equilibrate to room temperature for 15 mins. Bright-Glo (Promega) Luciferase reagent was then added at 100 µL/well. After a 2-minute incubation period at room temperature, luminescence was measured using a PerkinElmer plate reader. Cytotoxicity was performed using the same cells but without virus and monitored using CellTiter-Glo according to the manufacturer’s instructions.

### Time-of-drug addition assay

Time-of-drug addition assay was performed in Vero CCL81 cells. Cells were plated at 1×10^4^ cells/well in a 96-well plate and infected with SARS2-S. Compounds SR-914 (10 µM) and E64D (20 µM) were added to the wells at different time points post-infection. The infection proceeded for 48 hours. Bright-Glo (Promega) Luciferase reagent was used to quantify viral replication.

### ACE2 and TMPRSS2 mRNA expression

Total RNA was extracted using the RNA extraction kit (Qiagen) following the manufacturers instruction. Contaminating DNA was removed using Turbo-DNAse kit (Ambion). cDNA was synthesized using Sensifast (Bioline) following the manufacturer’s instructions. Real time qPCR was performed with an aliquot of cDNA as template, using Sensifast SYBR green (Bioline) in a 20 μL reaction according to the manufacturer’s instructions. The same validated primer sequences of GAPDH, ACE2 and TMPRSS2 were implemented as previously described^25^.

### Live virus inhibition assays

The cytopathic effect (CPE) assay was determined using Vero E6 cells expressing different levels of ACE2 cells and seeded at 12,000 cells/well in 96-well plates. The next day, compounds diluted in cell culture media were added. After 2 hours of incubation, SARS-CoV-2 (USA-WA1/2020 isolate; BEI resource) diluted at 600 pfu/15 µL was added to each well. After 72-hour incubation period, cell viability (protection from virus-induced CPE) was measured with CellTiter-Glo. Cytotoxicity was similarly measured, with culture media replacing the virus.

A virus titer reduction assay was performed as followed. Twelve-well plates with 100,000 Vero E6 cells grown overnight were infected with virus at a MOI of 0.05 diluted in the cell culture medium. Virus and cells were incubated together at 37°C for one hour to allow for sufficient adsorption. The unabsorbed virus was washed away with 1 mL of PBS once, and the wells were replenished with cell culture media with compound or DMSO (0.25% vol/vol, final). After 24 hours, supernatant and cells were harvested for further analysis. The progeny viruses in the supernatants were enumerated using the plaque assay with Avicel as described below. Vero E6 cells grown to confluence in 24-well plates were infected with 167 μL of serially diluted virus samples for one hour at 37°C and 5% CO_2_. Wells were washed with 1XPBS and replenished with virus infection medium and 0.8% Avicel (DuPoint). Three days after infection, viral infection centers were visualized with crystal violet staining (0.2% crystal violet, 4% paraformaldehyde and 10% ethanol).

### Luciferase complementation assays with 3CLpro and PLpro

The PLpro assay was run per the published methods^26^. The 3CLpro assay followed the exact same procedure albeit utilized the 3CLpro vector.

### Molecular modeling

The crystal structure of SARS-CoV-2 RBD bound with ACE2 (PDB ID: 6M0J) was imported to Schrodinger Maestro. The ACE2 receptor component of the crystal structure was removed and the RBD preprocessed at a pH of 7.4 +/- 0.2, using the protein preparation wizard within Schrodinger Maestro. A restrained minimization was performed utilizing the OPLS3e forcefield with heavy atoms converging to 0.30 angstroms. A receptor grid for docking experiments was generated with the receptor grid generation workflow by centering the grid on the relevant RBD residues that have been noted to be critical for stable interaction with ACE2 (K417, G446, Y449, Y453, L455, F456, A475, F486, N487, Y489, Q493, G496, Q498, T500, N501, G502, Y505). Using the virtual screening workflow, the ligands to be docked were settled, prepared at a pH of 7.4 +/- 0.2, and then docked with the XP (high precision) docking protocol.

To construct the United Kingdom (UK) and the South African (SA) SARS-CoV-2 RBD mutants, the appropriate point mutations were introduced to the previous structure (K417N, E484K, N501Y) and similar protocol to the wild type was performed. As for the D614G SARS-CoV-2 RBD mutant, the crystal structure of SARS-CoV-2 D614G 3 RBD down S Protein Trimer, without the P986-P987 stabilizing mutations, (PDB ID: 7KDK), was imported to Schrodinger Maestro. The 7KDK crystal structure was lacking loops and sidechains in the RBD that were added by using the Prime workflow of Schrodinger Maestro prior docking. The D614G and N501Y RBD mutant models were also derived from the 7KDK structure.

### Statistical analysis

P values were calculated using one-way or two-way analysis of variance (ANOVA) followed by a Tukey’s or Dunnett’s post hoc test. P values of <0.05 or <0.01 were considered statistically significant. Statistical analysis was performed using GraphPad Prism software (San Diego, CA, USA).

## RESULTS

### I. Screening data

#### 1. ReFRAME library Screening

The HTS campaign was initiated by primary screening against the ReFRAME library of 13,136 compounds at a single concentration of 10 µM, in singlicate (**Fig.1C**). Raw assay data was imported into Scripps’ corporate database to facilitate application of compound heredity to test wells and for statistical analysis. The primary data was then used to calculate a Z-score value, which showed a very reasonable distribution demonstrating the reliability of the assay (**Fig.1D**). The assay performance was robust with an average Z’ of 0.65 ± 0.064 and an average S:B of 146.0 ± 10.0 (n=12 plates, **Fig.1G**). As part of all HTS campaigns, we interleave multiple plates with 1280 wells each treated with vehicle only (DMSO plates) to ensure false positive and negative hit rates are negligible as well as to facilitate hit cutoff determination. Using data obtained from these plates three values were calculated: (1) the average value for all DMSO wells; (2) the SD value for the same set of data wells in expression 1; and the sum of (1) and 3 times of (2) was used as cutoff. Any compound that exhibited greater percent activation than the cutoff parameter was declared active. Using these criteria, the hit-cutoff was determined to be 55.7% activation, which yielded 990 compounds that exceeded that value (“hit”, **Fig.1C**).

To select compounds specifically targeting SARS2-S entry, we compared the activity of the 990 hits against other COVID-19 directed screens completed at Scripps. These screens were done using the same libraries and were tested against SARS-CoV-2 3CL protease (3CLpro) and SARS-CoV-2 papain-like protease (PLpro)^26^. The chymotrypsin-like 3CLpro cleaves the polyprotein at 11 distinct sites, while the PLpro cleaves the polyprotein at 3 sites and cleaves ubiquitin and ISG15 form target proteins for innate immune evasion^27^. **Fig.1F** shows the Venn diagram analysis of hits from SARS2-S entry, 3CLpro and PLpro. The analysis identified 419 compounds specific to SARS2-S entry. A confirmation screen was then run with these selected compounds, in triplicate, at 10 µM (**Fig.1C**). The SARS2-S entry confirmation assay yielded an average Z’ of 0.64 ± 0.025 and a S:B of 158.3 ± 2.5 (**Fig.1G**). Using a hit cutoff of 43.4 % inhibition derived from the average activity and 3 times the standard deviation of all DMSO treated (vehicle) wells, we confirmed the activity of 306 hits (73.0% hit rate; **Fig.1C**). In parallel, we assessed the cytotoxicity of the compounds with a live dead readout incorporating CellTiter-Glo detection reagent. This assay yielded an average Z’ of 0.96 ± 0.01 and a S:B of 141.9 ± 1.7 (**Fig.1G**). 227 hits were selected (54.2 %) using a cutoff of 12.0 % inhibition; and upon merging entry and cytotoxicity data, 108 compounds were found to inhibit SARS2-S entry without being overtly cytotoxic (**Fig.1C**). Next, these selected compounds were titrated using 10-point dose-response titrations (3-fold dilutions), in triplicate. The SARS2-S entry titration assay performance was consistent with an average Z’ of 0.60 ± 0.035 and a S:B of 157.0 ± 9.5. (**Fig.1G**). Of the 108 compounds tested, 74 compounds demonstrated nominal potency (IC_50_ < 10 µM) in the SARS2-S entry assay and were considered active. The cytotoxicity counterscreen identified 6 compounds with a CC_50_ < 10 µM, and the VSV-G counterscreen identified 42 compounds with an IC_50_ < 10 µM (**Fig.1C**). At the conclusion of the HTS phase, we found 6 small molecules of interest from the ReFRAME collection with a therapeutic index (TI=CC_50_/IC_50_) higher than 15 (**Table.S1**). However, upon further inspection, we learned that these 6 ReFRAME compounds including their analogs had recently been identified by other groups when tested in whole virus CPE assays^16, 28, 29^, which confirmed the robustness of our assay but diminished our interest in pursuing these compounds further. As such, we focused our attention on the hits identified from the other collections.

#### 2. TargetMol, Pathogen Box and Cathepsin L libraries screening

The aforementioned method was also used to screen a total of ~2K compounds from the TargetMol, Pathogen Box and Cathepsin L libraries (**Fig.1E and G**). In this instance, compounds were tested at a single concentration of 4 µM in triplicate. The assay performance was excellent (TargetMol: Z’ of 0.63 ± 0.08 and a S:B of 150.06 ± 7.76; Pathogen Box: Z’ of 0.62 ± 0.05 and a S:B of 134.73 ± 6.17; Cathepsin L: Z’ of 0.70 ± 0.05 and a S:B of 140.64 ± 5.45) and a cutoff of 54.2% inhibition, similarly derived, yielded 91 hits from the TargetMol library, 15 from the Pathogen Box and 5 hits from the Cathepsin L library (**Fig.1E**). From a total of 111 hits, all but two compounds were available for titration assays. As with the ReFRAME library, all 109 compounds from these libraries were tested in 10-point dose-response titrations (3-fold dilutions) in triplicate. All compounds had IC_50_ values < 10 µM in this assay. The cytotoxicity counterscreen identified 61 active compounds, while the VSV-G counterscreen titration assay identified 90 compounds with an IC_50_ < 10 µM (**Fig.1E**).

Out of the 91 hits, 11 showed a TI higher than 20 and a least ~10-fold higher sensitivity to SARS2-S as compared to VSV-G (**Table 1**). Interestingly, SR-914, SR-372 and SR728 had a TI and specificity toward SARS2-S higher than achieved by any ReFRAME hit. As compared to the best ReFRAME hit, SR-806 (VBY-825), SR-914 and SR-372 showed 7.1 and 3.5-fold higher potency, and a specificity higher than 7.1 and 3-fold, respectively (**Table 1 and S1**).

#### 3. Enrichment of inhibitors toward known targets (**Fig.2A**)

Analysis of the known targets of the hit compounds from all libraries revealed that 40% of the small molecules identified had previously been shown to target cysteine proteases that are known to be involved in SARS-CoV-2 entry^30–34^. Another 7% of the selected hits target Lanosterol 14 alpha-demethylase, which is involved in sterol biosynthesis and glucosylceramide synthase, which is important for glucosylceramide expression. Sterol and glucosylceramide are components of lipid rafts, which influence entry for a variety of viruses^35^. In particular, cholesterol has been proposed to be important in SARS-CoV-2 lipid dependent entry^36^. The implication of glucosylceramide in SARS-CoV-2 entry was also previously suggested, using multiple glucosylceramide synthase inhibitors against SARS-CoV-2^29^. The depletion of the glucosylceramide from the cell membrane has however not yet been investigated.

**Fig.2.**
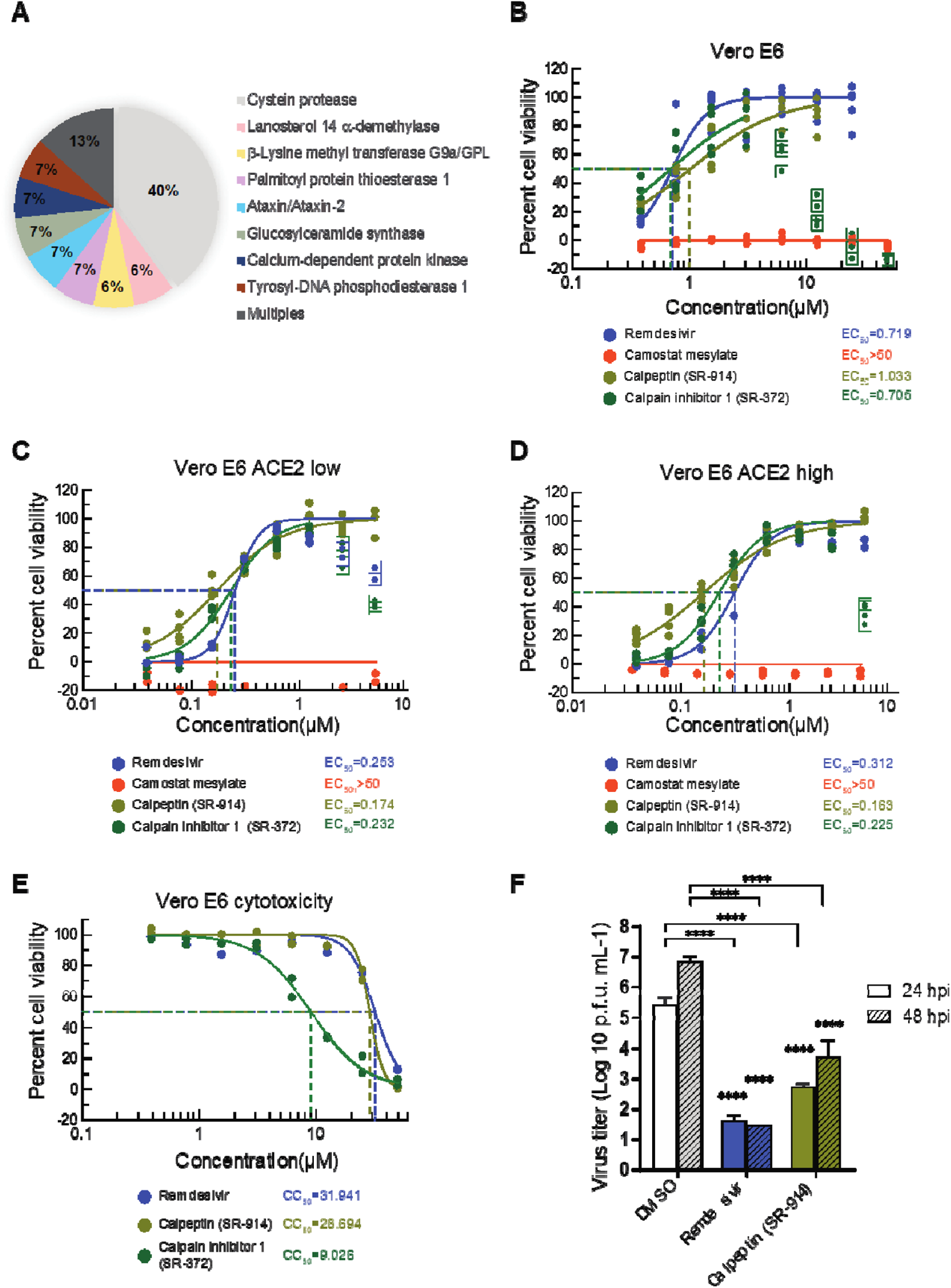
Targets of the selected compounds and SARS-CoV-2 wild type infection. A. Description of the targets of the different hits from all the studied libraries. **B. Antiviral activity of the 2 best hits in the SARS-CoV2-induced CPE assay.** Vero E6 cell treated with test compounds for two hours was infected with SARS-CoV-2 virus at an MOI of 0.05, then incubated for three days in the presence of compound. Cell viability (protection from virus-induced CPE) was measured with CellTiter-Glo. **C and D.** Antiviral effect was measured with a subset of Vero E6 cells expressing a low (**C**) and high (**D**) level of ACE2. **E. Cytotoxicity of selected compounds in Vero E6 cells.** Cytotoxicity was tested in the same conditions with cell culture media instead of the virus. **F. Virus yield reduction activity of selected compounds.** Vero E6 cells infected with SARS-CoV-2 at an MOI of 0.05 were cultured in the presence of test compound (5 µM) and the supernatant was harvested after 24 and 48 hours of incubation. The Progeny virus was enumerated with a plaque assay using an Avicel overlay in fresh Vero E6 cells. N=3 experiments were performed for infectivity assays and n=2 for the cytotoxicity assays. **** P< 0.0001, Two-way ANOVA with Dunnett’s multiple comparisons test against DMSO control.

Inhibitors of calcium dependent protein kinase were also among the hits we identified. Coronaviruses are reported to use calcium ions during entry into host cells^37, 38^, and experimental depletion of calcium seems to reduce viral entry^38^. It has been reported that SARS-CoV-1 S protein stimulates cyclooxygenase-2 (COX-2) expression *via* both calcium-dependent and calcium-independent protein kinase C pathways^39^. This subsequently results in activation of inflammatory pathways, eventually leading to adult respiratory distress syndrome (ARDS) in some individuals^40^. Individuals with COVID-19 experience marked increase in COX metabolites associated with bronchoconstriction and recruitment and activation of inflammatory cells and platelets^41^. The use of COX-2 inhibitors early on in the disease has been recommended^42^, suggesting a new role of SARS-CoV-2 S protein in inflammation, not previously described.

Finally, we also identified small molecules targeting the Palmitoyl-protein thioesterase 1, an autophagy modulator and the target of chloroquine derivatives ^43^, along with a few molecules that target unknown proteins for viral entry, namely tyrosyl-DNA phosphodiesterase 1, ataxin, and beta-Lysine methyl transferase G9a/GPL. These may merit further investigation.

### II. Lead activity against wild type SARS-CoV-2

Compounds with a therapeutic index (TI) greater than 100 (SR-914, SR-372, SR-510, SR-728 and SR-687) were further investigated for their ability to inhibit live SARS-CoV-2 replication by assessing their *in vitro* CPE. We infected Vero E6 cells with the original SARS-CoV-2 isolate, USA-WA1/2020, in the presence of increasing concentrations of compounds or DMSO control (**Table.S2, Fig.2B**), while monitoring compounds cytotoxicity in parallel (**Table.S2, Fig.2E**). SR-914 (Calpeptin) and SR-372 (Calpain inhibitor I) showed a TI > 10, with SR-914 having a similar potency as the nucleoside analog Remdesivir^8, 44, 45^ (**Fig.2B**). SR-510, SR-728 and SR-687 were inactive against wild type SARS-CoV-2 (**Table.S2**). Similarly, Camostat mesylate, a previously reported SARS-CoV-2 fusion entry inhibitor, did not shown any activity. Next, we tested SR-914 and SR-372 in Vero E6 that presented either low (23%) or high (88%) levels of surface of ACE2 receptor (**Fig.S2A**). This experiment revealed that the differential ACE2 expression levels did not substantially impact compound efficacy. In Vero cells with either low or high ACE2 expression, SR-914 showed an EC_50_=0.174 and 0.163 µM and SR-372 EC_50_=0.232 and 0.225 µM; respectively. Remdesivir displayed a similar EC_50_ in cells with high ACE2 expression, while Camostat mesylate was again inactive (**Fig.2C and D**). The activity of the best performing compound, SR-914, was further confirmed in plaque assays of wild type SARS-CoV-2 infection of Vero E6 cells at 24 h and 48 h post-treatment (**Fig.2F**). A similar shift of the potency of both SR-914 and SR-372, as well as the Cathepsin inhibitor control, E64d, was observed using SARS2-S entry assay in Vero CCL81 cells, which express lower levels of ACE2 receptor as compared with HEK293T-ACE2 cells (**Table.1 and S1, Fig.S1 and S2B**). Altogether, these results suggest a specific inhibitory mechanism of SR-914 and SR-372, dependent on ACE2-mediated entry.

**Table.1.**
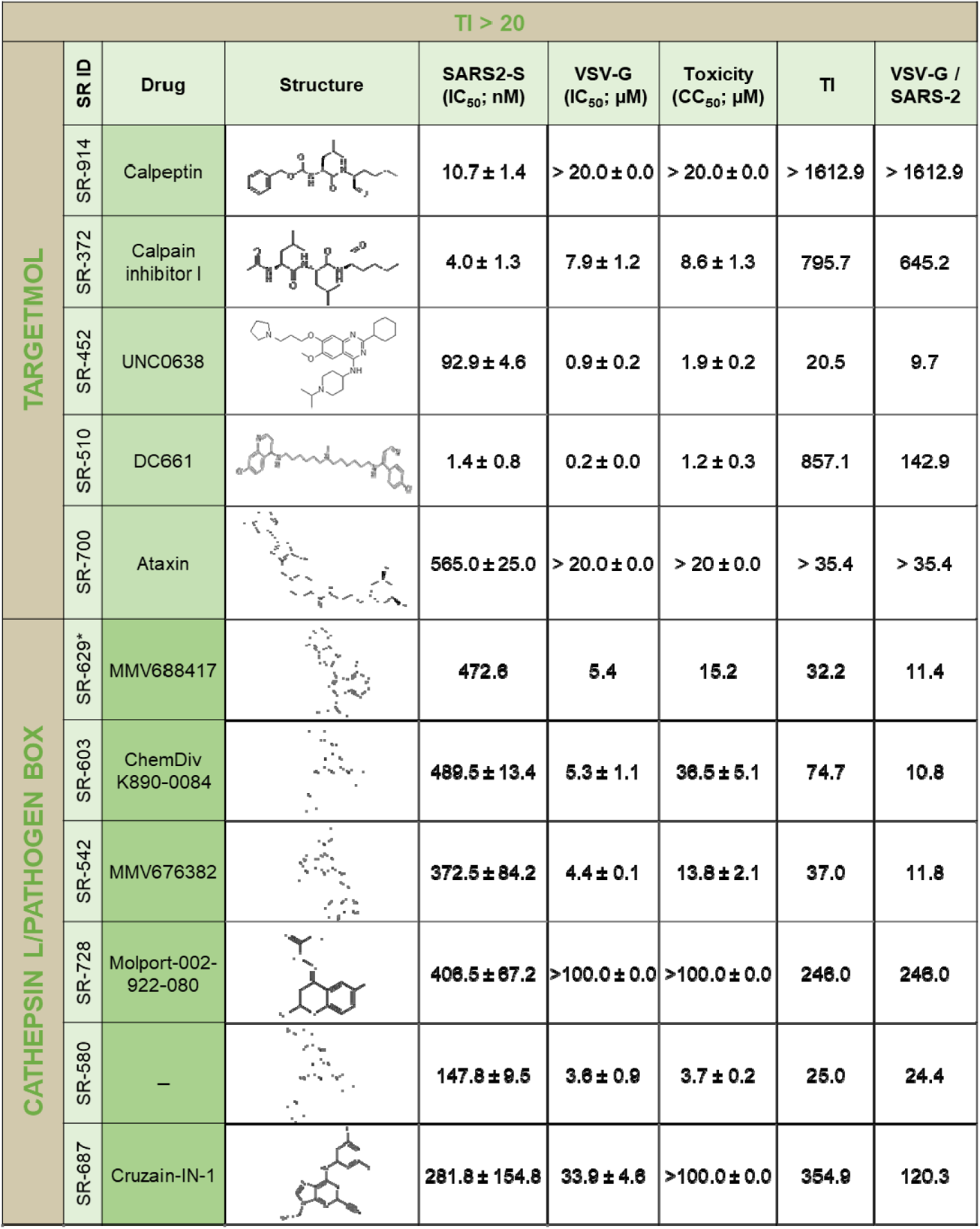
Summary of the selected cathepsin L, Pathogen box and TargetMol compounds in this study. Activity of the selected compounds against the different MLV pseudotyped viruses in HEK293-ACE2 cells and their respective cytoxicity. Values for SARS2-S, VSV-G and toxicity are mean ± SEM of 2-4 independent experiments. TI: therapeutic index. * n=1.

| TI > 20 |  |  |  |  |  |  |  |  |
| --- | --- | --- | --- | --- | --- | --- | --- | --- |
| TARGETMOL | SR ID | Drug | Structure | SARS2-S<br>(IC<sub>50</sub>; nM) | VSV-G<br>(IC<sub>50</sub>; μM) | Toxicity<br>(CC<sub>50</sub>; μM) | TI | VSV-G /<br>SARS-2 |
|  | SR-914 | Calpeptin |  | 10.7 ± 1.4 | > 20.0 ± 0.0 | > 20.0 ± 0.0 | > 1612.9 | > 1612.9 |
|  | SR-372 | Calpain inhibitor I |  | 4.0 ± 1.3 | 7.9 ± 1.2 | 8.6 ± 1.3 | 795.7 | 645.2 |
|  | SR-452 | UNC0638 |  | 92.9 ± 4.6 | 0.9 ± 0.2 | 1.9 ± 0.2 | 20.5 | 9.7 |
|  | SR-510 | DC661 |  | 1.4 ± 0.8 | 0.2 ± 0.0 | 1.2 ± 0.3 | 857.1 | 142.9 |
|  | SR-700 | Ataxin |  | 565.0 ± 25.0 | > 20.0 ± 0.0 | > 20 ± 0.0 | > 35.4 | > 35.4 |
| CATHEPSIN L/PATHOGEN BOX | SR-629* | MMV688417 |  | 472.6 | 5.4 | 15.2 | 32.2 | 11.4 |
|  | SR-603 | ChemDiv<br>K890-0084 |  | 489.5 ± 13.4 | 5.3 ± 1.1 | 36.5 ± 5.1 | 74.7 | 10.8 |
|  | SR-542 | MMV676382 |  | 372.5 ± 84.2 | 4.4 ± 0.1 | 13.8 ± 2.1 | 37.0 | 11.8 |
|  | SR-728 | Molport-002-922-080 |  | 406.5 ± 67.2 | >100.0 ± 0.0 | >100.0 ± 0.0 | 246.0 | 246.0 |
|  | SR-580 | — |  | 147.8 ± 9.5 | 3.6 ± 0.9 | 3.7 ± 0.2 | 25.0 | 24.4 |
|  | SR-687 | Cruzain-IN-1 |  | 281.8 ± 154.8 | 33.9 ± 4.6 | >100.0 ± 0.0 | 354.9 | 120.3 |

### III. SR-914 characterization

Since SR-914 and SR-372 are analogs (**Table.1**) and SR-914 showed improved TI, we emphasized studies with SR-914.

#### 1. Activity of SR-914 against the cell surface entry pathway

As SR-914 significantly inhibited the endosomal pathway, we verified its activity against the second SARS-CoV-2 reported entry pathway, the cell surface entry pathway, where the S protein is activated by TMPRSS2 at (or close to) the cell surface, resulting in fusion of the viral and plasma cell membranes^46–48^. We used HEK293T stably expressing ACE2 and TMPRSS2 cells (**Fig.3A, S2B**) in SARS2-S and VSV-G entry assays. SR-914 inhibited SARS2-S with an IC_50_ of 10.93 ± 0.78 nM and VSV-G with an IC_50_ > 20 µM. These results are similar to the activity of the compound in HEK293T-ACE2 cells devoid of TMPRSS2 (**Table.1, Fig.S2B**), suggesting the ability of SR-914 to inhibit SARS2-S entry independently of TMPRSS2, and likely the ability of the compounds to inhibit both reported entry pathways for SARS-CoV-2.

**Fig.3.**
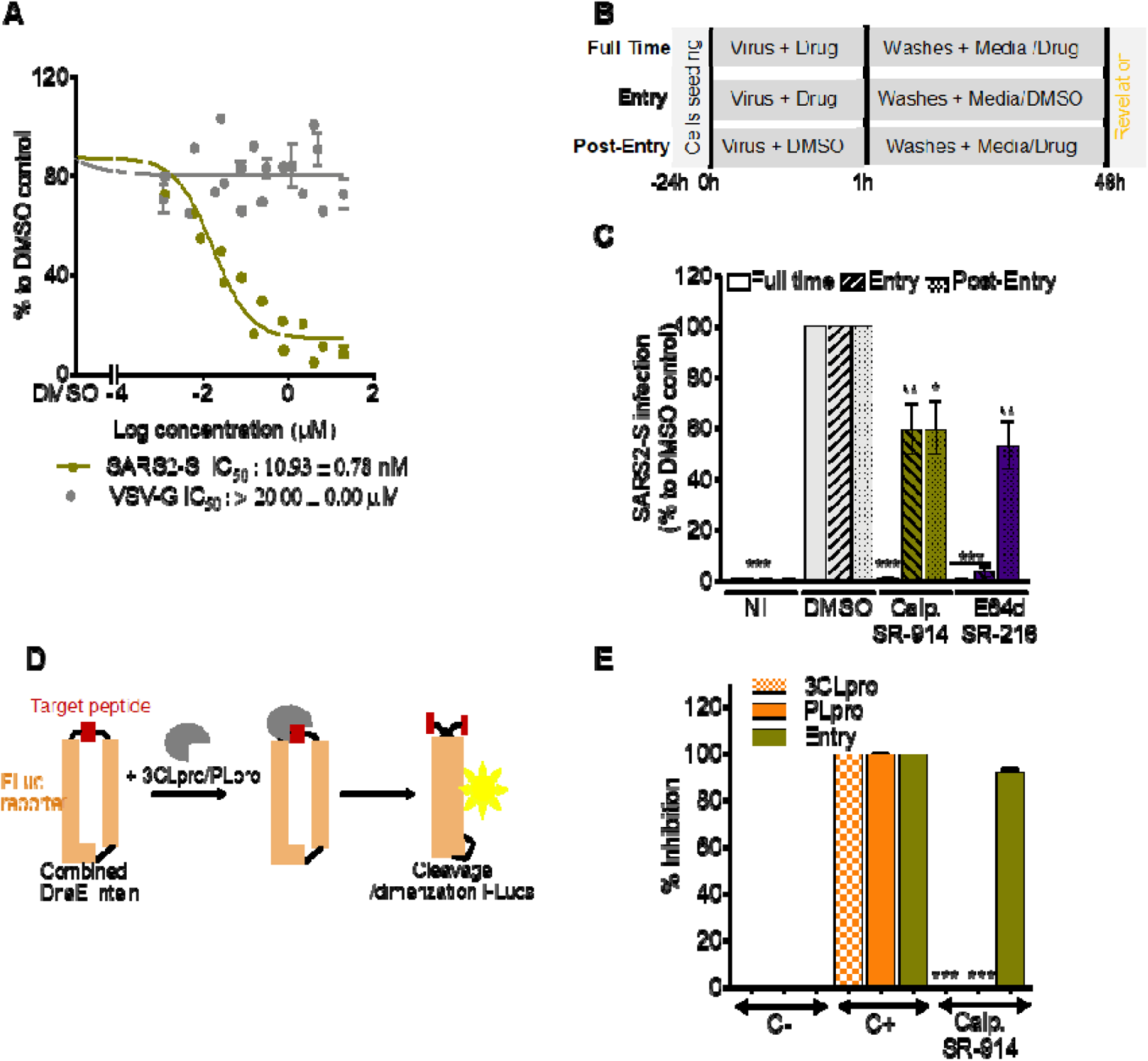
SR-914 “Calpeptin” specifically blocks SARS-CoV entry. A. Its activity against SARS2-S in HEK293T-ACE2-TMPRSS2 cells. Cells were incubated with different concentrations of drugs, then infected with SARS2-S or VSV-G. Luciferase was measured 48 hours later, using Bright-Glo. Shown is the mean ± SEM of n=2 to 4 independent experiments. **B. Time of drug addition experiment schematic.** Infection was performed for 1 h with or without drugs, Vero CCL81 cells were then washed, and fresh media was added with or without drugs**. C. Time of drug addition experiment result.** SR-914 was used at 10 µM. E64d at 20 µM. Calp. : Calpeptin = SR-914. NI: not infected. Shown is the mean ± SEM of 4 to 6 independent experiments. **D. Luciferase complementation assay schematic.** The reporter consists of a split Firefly luciferase protein connected by a cleavable peptide for the tested protease. Upon cleavage of the peptide, the luciferase protein undergoes dimerization for an active state. DnaE intein helps in this dimerization. **E. Its activity against SARS2-S Entry, 3CLpro and PLpro.** C-: negative control. C+; positive control. Shown is the mean ± SD of 3 independent experiments. One-way ANOVA followed by Tukey’s post-test were used for statistical comparisons. *, P < 0.01; **, P < 0.001; ***, P < 0.0001.

#### 2. Mechanism of action of SR-914

To further confirm the stage of SARS2-S entry targeted by SR-914, we conducted time-of-drug addition experiments, using Vero CCL81 cells. Compounds were either incubated with cells during the virus entry process (entry), post-entry (post-entry) or during the entire infection period (full-time), and luciferase expression was quantified 48 hours later (**Fig.3B**). As anticipated, total inhibition was observed when SR-914 was present during the “full-time” condition. Interestingly, SR-914 blocked equally well during both entry and post-entry stages (**Fig.3C**). In the case of the “entry” condition, compound was only present during the first hour prior to removal of excess virus by washing. In the “post-entry” condition, compound was only present after the washing step, where the virus had already permeated the cell membrane. Interestingly, VBY-825, which is a cathepsin inhibitor from the ReFRAME library, previously shown to be an entry inhibitor, presented a similar pattern as SR-914^49^. However, E64d showed a better potency at the entry than post-entry (**Fig.3C**). This result suggests a dual mechanism of action for SR-914 at the entry and post-entry steps.

#### 3. Specificity of SR-914

SR-914 is a cysteine protease inhibitor (Calpain I and II, Cathepsin L and K). To verify its specificity toward the entry of SARS2, we tested its activity against the SARS-CoV-2 non-structural proteases 3CLpro and PLpro in a luciferase complementation reporter assay, which incorporates the unique peptide cleavage site specific for each of the proteases (**Fig.3D**). In this sense, inhibitors of these assays would diminish the luminescence response generated upon implementation of the firefly luciferase substrate. SR-914 showed no activity against 3CLpro and PLpro (**Fig.3E**), further supporting a specific role for SR-914 in the inhibition of the early events involved in SARS-CoV-2 entry.

#### 4. Activity of SR-914 against viruses pseudotyped with the S protein from SARS-CoV-1 and highly virulent emerging strains of SARS-CoV-2

The breath of SR-914 activity was first investigated in entry assays in HEK293T-ACE2 cells of MLV reporter viruses pseudotyped with the S proteins of SARS-1 (**Fig.4A**). SR-914 showed potent activity against SARS1-S protein with an IC_50_ of 77.18 ± 4.69 nM. However, it was found to be ~8 fold less potent than it is against SARS2-S (**Fig.4A and Table 1**). This lower activity may be derived from the fact that only 68% similarity is shared between the two S proteins^50, 51^, suggesting specificity of SR-914 to the SARS-CoV-2 entry step.

**Fig.4.**
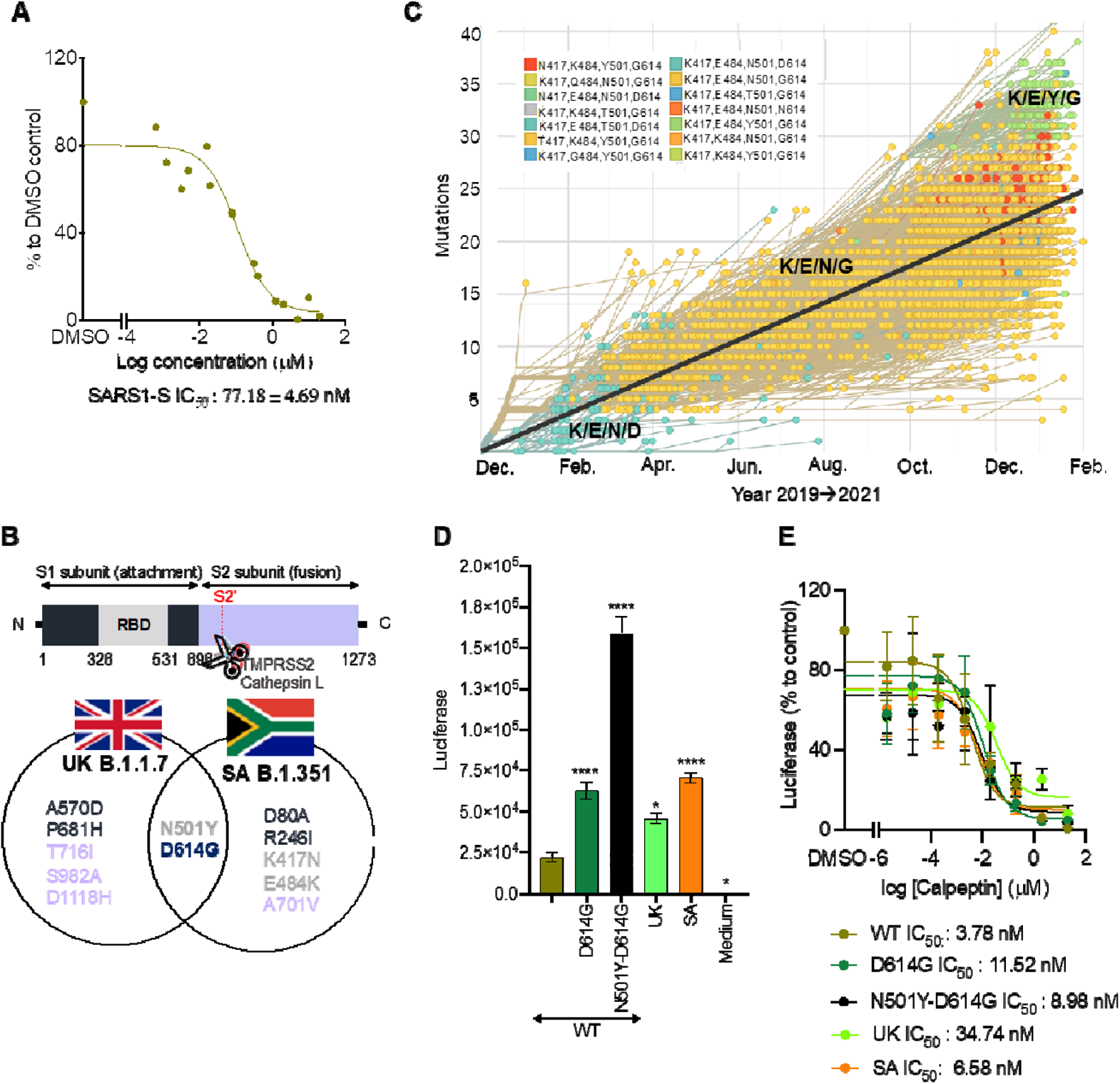
Breath of activity of Calpeptin against various SARS-CoVs. A. Its activity against SARS1-S in HEK293T-ACE2 cells. HEK293T-ACE2 cells were incubated with different concentrations of Calpeptin, then infected with SARS1-S. Luciferase was measured 48 hours later, using Bright-Glo. Shown is the mean ± SEM of n=2 independent experiments. **B. Schematic of the substituted residues in the S protein of the highest threat of SARS-CoV-2 strains. C. Evolution of the S protein residues at the position 417, 484, 501 and 614 from 2019 to February 2021.** Modified figure from https://nextstrain.org/ncov/global?branchLabel=none&c=gt-S_417,484,501,614&l=clock**. D. Activity of the new emergent variants.** HEK293T-ACE2 cells were infected with different mutants of SARS2-S. The day after, a medium change was performed. Luciferase was measured 48 hours later, using Bright-Glo. Shown is the mean ± SEM of n=3 independent experiments. WT: wild type, SA: South Africa, UK: United Kingdom. **E. Activity of Calpeptin activity against crucial mutations present in the S protein of the new emergent strains.** Similar experiment than D but Calpeptin was added during infection and after medium change. Shown is the mean ± SEM of n=2-5 independent experiments. Two-way ANOVA followed by Dunnett’s post-test were used for statistical comparisons. *, P < 0.01; **, P < 0.001; ***, P < 0.0001.

New SARS-CoV-2 variants have been emerging worldwide (**Fig.4B and C**). The United Kingdom (UK) variant, known as B.1.1.7, has large number of variations but noteworthy is the N501Y-D614G mutation located in the S protein, which has been associated with increased risk of transmissibility^10, 52^. The South Africa (SA) variant, B.1.351 also has these 2 substitutions and additional mutations at residues K417N and E484K that weaken the neutralization by current vaccines^11, 52^. Residues 417, 484 and 501 are located in the crucial 438-506 region of the RBD involved in the binding of the S protein with the ACE2 receptor^53^. Although the mutation 614 is far from this region, it was shown to affect their interaction^54^. All these variants have now been detected in numerous countries, including the United States. The D614G mutation variants overtook the wildtype SARS-CoV-2 over the year of 2020 but importantly in 2021 an increased number of populations are affected by N501Y-D614G variants (**Fig.4C**). Delta, or B.1.617.2, is now circulating in 98% of the countries world wide and is considerably more transmissible (~2 fold) than the first Wuhan strain. It has ~10 mutations in the S glycoprotein, with the four of them associated to higher virulence (L452R, T478K, D614G and P681R ^55^). These may support the increase in vaccine breakthrough recently observed with this variant. In fact, the WHO currently regards the Delta variant as the fittest and fastest variant so far. Notably, at the time that most of the work associated with this manuscript was done, the delta variant didn’t exist, again alluding to the expediency with which incredibly infectious variants arise.

We thus investigated the activity of SR-914 in entry assays in HEK293T-ACE2 cells infected with pseudotyped viruses expressing the S protein mutated in residues D614G, N501Y-D614G or presenting the mutations from UK and SA variants (**Fig.4D and E**). Pseudotyped viruses were used at the same TCID_50_.

In the absence of Calpeptin, all construct presenting inclusive of the mutation D614G showed a significant increase of luciferase expression, when used at equivalent MOI, compared to Wild Type (WT, **Fig.4D**). This increased activity related to the mutation D614G was previously shown by Choe’s group^54^. Interestingly, the combination of this mutation with N501Y further amplified the luciferase signal and confirms the importance of this mutation in the increased spread of the virus (**Fig.4D**). To our knowledge this has never been demonstrated *in vitro*. Only a recent computational analysis of mutational strains^56^ described N501Y as a more prone to disease compared to D614G. The mutation of D614G for secondary structure prediction shows no changes in secondary structure, while remaining in the coil region, whereas the mutation N501Y changes from coil structure to extended strand. This results in a higher affinity to human ACE2 protein compared to D614G based on this docking study. Remarkably, N501Y-D614G mutations present in UK and SA strains did not induce the same increase of activity (**Fig.4D**), suggesting that the other mutations of both strains may have counteract the activity of the double mutations.

Treatment with Calpeptin (SR-914) reduces the infectivity of all the mutants, in the nanomolar range, with marginally different potency. Calpeptin inhibits the constructs in the following order:

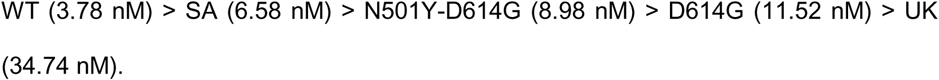

As we mentioned earlier, we hypothesized that Calpeptin may have a dual mechanism of action that inhibit virus entry and post entry steps. To visualize how calpeptin inhibits entry steps, we performed molecular docking of the compound SR-914 to wildtype RBD in the context of the ACE2 receptor (**Fig.S3**). Accordingly, *in silico* ligand docking studies revealed that Calpeptin binds to the WT RBD (PDB ID: 6M0J), with the highest affinity (−4.38 kcal/mol), by forming two hydrogen bonds with S494, a single hydrogen bond with Y453, a π-π stacking interaction with Y505, and a p-cation interaction with R403 (**Fig.S3D**). These residues are all critical to forming the intermolecular interaction with the ACE2 receptor^53^, and therefore viral entry. As expected, an inactive analog of Calpeptin showed poor binding to WT RBD (−3.46 kcal/mol, **Fig.S5A**). Similar to the cell-based-assay results, Calpeptin has the lowest affinity for RBD-UK (derived from PDB ID: 6MOJ, −3.68 kcal/mol, **Fig.S4C**). This lower affinity likely stems from the N501Y mutation within the RBD, which results in Calpeptin binding in an alternate orientation that is less conducive to forming intermolecular bonds. When docked to the RBD-UK, Calpeptin forms three intermolecular interactions compared to the five intermolecular interactions formed with WT RBD. The D614G, N501Y-D614G and SA mutants had intermediate affinities for Calpeptin, docking at −4.04, −4.35 and −4.29 kcal/mol, respectively (**Fig.S4A, B, D**). The SA variant contains the K417N, E484K and N501Y mutations. The K417N and E484K mutations are outside of the immediate binding site of Calpeptin; however, Calpeptin forms a hydrogen bond with Y501 of the SA variant spike protein. The ability of Calpeptin to form a hydrogen bond with Y501 of the SA variant but not Y501 of the UK variant is likely the result of structural rearrangements that alter the shape of the binding cleft and can be attributed to the two additional mutations present in the SA variant that are distal from the Calpeptin binding cleft. Calpeptin docked to the D614G and N501Y-D614G RBD mutants in a similar orientation to that of the WT RBD. The N501Y-D614G RBD showed weaker affinity for Calpeptin relative to the D614G mutant through *in vitro* cell-based assay but the *in silico* ligand docking GlideScore is similar to WT. Consequently, the double mutant is the only outlier in the linear correlation plot (**Fig.S5B**). Both mutants form hydrogen bonds with S494 and G496, though the introduction of the N501Y mutation, which allows the formation of a hydrogen bond and an additional π-π stacking interaction bond between Calpeptin and the Y501 residue. Additionally, the introduction of the N501Y mutation created an additional π-π stacking bond between Calpeptin and Y505, likely as a result of a conformational perturbation originating from the Y501 mutation.

Mutant RBD models derived from PDB ID: 7KDK contained some missing residues within the RBD that were assigned from the protein peptide sequence using Prime in Schrodinger Maestro. Notably, the missing residues in the conformations are not derived from a crystal structure, and hence the models may stray from the actual structure of the mutant spike proteins. However, plotting the GlideScores obtained through *in silico* docking against the logarithmic IC_50_ of the ligand obtained in cell-based assays resulted in a strong linear correlation with a R^2^ of 0.878, validating the models created.

In summary, in this study, calpeptin showed inhibitory activity specifically to SARS2 entry as well as post-entry step. This strongly suggests calpeptin may a dual mechanism. For SARS entry inhibition, we hypothesized a direct binding between the RBD and Calpeptin that affects the S-ACE2 interaction. In support of this hypothesize, we point to the results of the time of drug addition experiments, the difference in activity of calpeptin against the various mutants of SARS2-S or SARS1-S, and the molecular modeling data. It is also quite plausible that Calpeptin inhibits SARs entry visa via inhibition of the host cell dependent pathway involving cathepsin L1. Calpain inhibitors such as Calpeptin are cell permeable protease inhibitors which have also been shown to inhibit the main viral protease (MPRO) albeit very weakly in the double digit micromolar range ^57 58^. Thus, its potent activity against the novel variants is a welcome feature given that these variants are rapidly overtaking the original, rapidly spreading, and escaping from current vaccines. For instance, J&J’s vaccine only inhibits the SA strain by 57%^59, 60^.

## Discussion

Our robust and reliable HTS campaign of ~15,000 clinically relevant compounds targeting SARS-CoV-2 entry identified SR-914 “Calpeptin”, which is a reversible semi-synthetic peptidomimetic aldehyde inhibitor of Calpain I, Calpain II, Cathepsin L and K^61, 62^ and now proven as a potent SARS-CoV-2 inhibitor in-vitro. In particular, Calpeptin inhibits USA-WA1/2020 isolate with a higher potency than the FDA approved drug Remdesivir in Vero E6 cells enriched in ACE2 receptors, and as shown through a dual temporal activity, allowing the inhibition of both SARS-CoV-2 entry pathways. Calpeptin has broad and specific activity against the novel highly virulent variants and SARS-CoV-1. We hypothesized calpeptin acts partially by binding to the RBD and preventing the binding to the ACE2 receptor, but may also prevent shedding at the entry step, as Calpeptin was previously shown to block the shedding of the prion proteins^63^. Additionally, Calpeptin may block S protein triggering calpain release from endoplasmic reticulum stress resulting in SARS-CoV-2 replication at the post-entry steps. This activity of the S protein was previously described for SARS-CoV-1 S protein^64^. The inhibition of SARS2-S by an inhibitor of a calcium dependent protein kinase pathway, SR-629 (**Fig.2A**), reinforces this hypothesis and provides SARS-CoV-2 S protein with a novel function. However, we do not exclude its anti-Cathepsin L activity at the post-entry step.

Importantly, Calpeptin potently blocked, in mice, aberrant calpain activities that display similar diagnoses to coronaviruses. In particular, Calpeptin inhibited calpain inducing lung related diseases such as inflammatory pulmonary, pulmonary fibrosis, ventilator-induced dysfunction of the diaphragm, asthma and anti-lung tumor^65–70^. It was also shown to be neuroprotective^71–74^ and to preserve the myocardial structure and function^75^. Crucially, Calpeptin was used in mice for about one month without any toxicity^71, 73^. These multiple and ubiquitous activities of Calpeptin could be very helpful in the severe state of the disease, where several organs are affected, and unfortunately poorly targeted by the FDA approved therapeutics. The use of calpain inhibitors are likely safe since several selective calpains inhibitors are currently in clinical trials for various diseases^58, 76–79^ and the compounds against SARS-CoV-2 are intended for use for acute disease control; where short-term toxicity and promiscuity may be better tolerated.

Taken together, our study demonstrates the success of a large-scale drug repurposing effort, confirming the outcomes of others with respect to hit-to-lead efficacy. We identified Calpeptin as a novel lead that appears to be a potent inhibitor of the early events in SARS-CoV-2 entry and confirmed its broad activity against whole virus and pseudotyped virus containing novel alarming coronavirus variants. Our future efforts will be focused on determining the precise mechanism of action, testing its activity in human tissues, and performing structure-activity relationship along with using it as a component of inhibitor cocktails for increased potency.

## Acknowledgements

We thank Lina DeLuca (Lead Identification, Scripps Florida) for compound management, Dr. Christopher Rader, and Dr. Haiyong Peng (Scripps Florida) for their advice and help.

## Supplemental tables and figures

**Table.S1.**
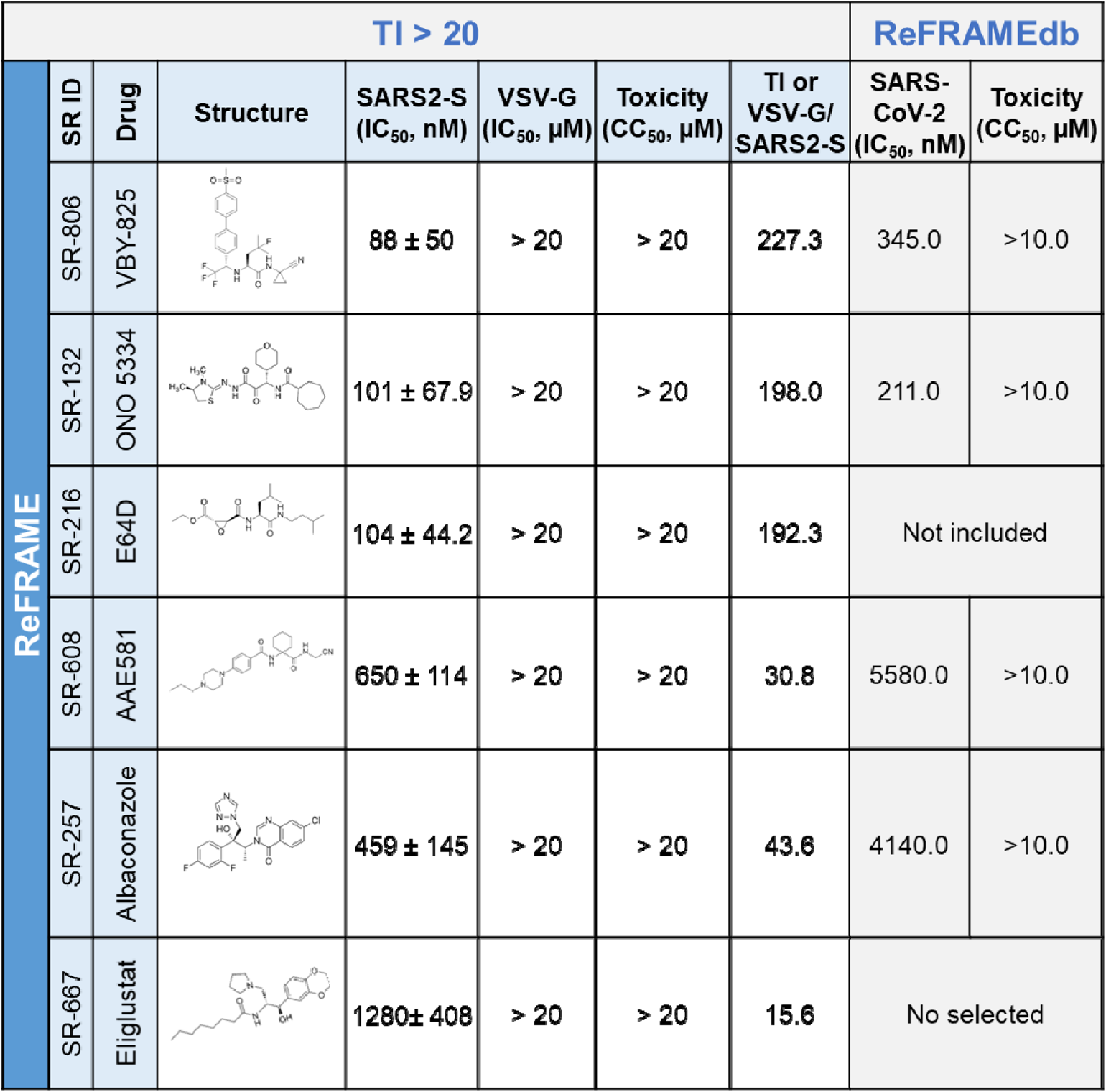
Summary of the ReFRAME compounds with a therapeutic index higher than 20. Activity of the selected compounds against the different MLV pseudotyped viruses in HEK293-ACE2 cells. Values for SARS2-S, VSV-G and toxicity are mean ± SD of 3 independent experiments. ReFRAMEdb: activity of the compounds was measured using the cytopathic effect (CPE) of the SARS CoV-2 virus infecting Vero E6 host cells. The viability of uninfected cells after exposure to hit compounds for 72 hours is measured to determine compounds cytotoxic effects. TI: therapeutic index.

| TI > 20 |  |  |  |  |  |  |  | ReFRAMEdb |  |
| --- | --- | --- | --- | --- | --- | --- | --- | --- | --- |
| ReFRAME | SR ID | Drug | Structure | SARS2-S<br>(IC<sub>50</sub>, nM) | VSV-G<br>(IC<sub>50</sub>, μM) | Toxicity<br>(CC<sub>50</sub>, μM) | TI or<br>VSV-G/<br>SARS2-S | SARS-CoV-2<br>(IC<sub>50</sub>, nM) | Toxicity<br>(CC<sub>50</sub>, μM) |
|  | SR-806 | VBY-825 |  | 88 ± 50 | > 20 | > 20 | 227.3 | 345.0 | >10.0 |
|  | SR-132 | ONO 5334 |  | 101 ± 67.9 | > 20 | > 20 | 198.0 | 211.0 | >10.0 |
|  | SR-216 | E64D |  | 104 ± 44.2 | > 20 | > 20 | 192.3 | Not included |  |
|  | SR-608 | AAE581 |  | 650 ± 114 | > 20 | > 20 | 30.8 | 5580.0 | >10.0 |
|  | SR-257 | Albaconazole |  | 459 ± 145 | > 20 | > 20 | 43.6 | 4140.0 | >10.0 |
|  | SR-667 | Eliglustat |  | 1280 ± 408 | > 20 | > 20 | 15.6 | No selected |  |

**Table.S2.**
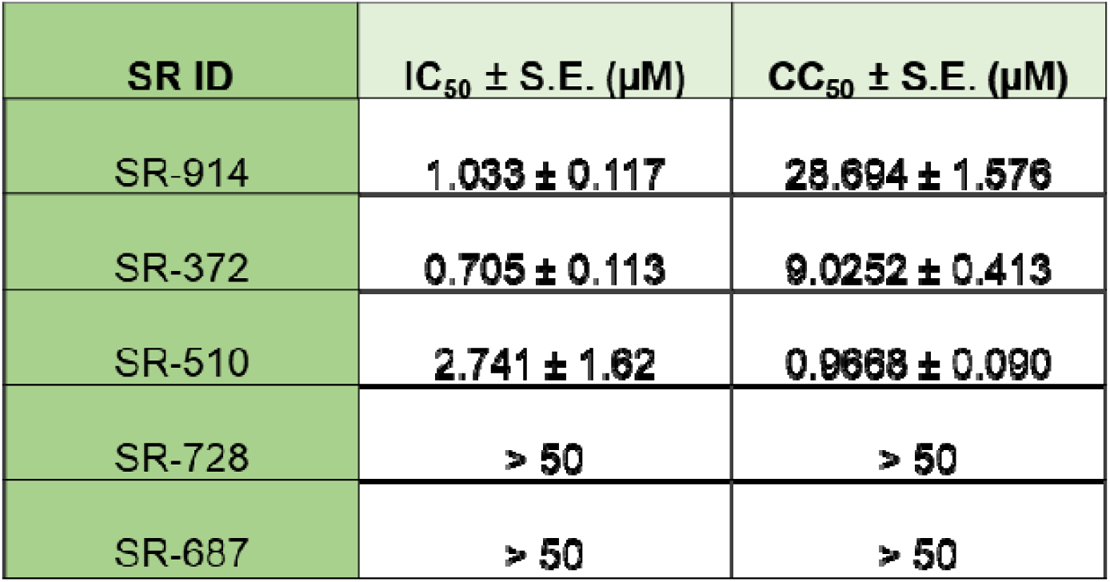
Anti-antiviral activity of the compounds with therapeutic index higher than 100. Vero E6 cell treated with test compounds for two hours was infected with SARS-CoV-2 virus at an MOI of 0.05, then incubated for three days in the presence of compound. Cell viability (protection from virus-induced CPE) was measured with CellTiter-Glo. Cytotoxicity was tested in the same conditions with cell culture media instead of the virus. IC_50_ and CC_50_ were calculated with the 4-parameter Logistic model (XLFit fit model 205) and the standard errors (S.E.) were shown.

| SR ID | IC<sub>50</sub> ± S.E. (μM) | CC<sub>50</sub> ± S.E. (μM) |
| --- | --- | --- |
| SR-914 | 1.033 ± 0.117 | 28.694 ± 1.576 |
| SR-372 | 0.705 ± 0.113 | 9.0252 ± 0.413 |
| SR-510 | 2.741 ± 1.62 | 0.9668 ± 0.090 |
| SR-728 | > 50 | > 50 |
| SR-687 | > 50 | > 50 |

**Fig.S1.**
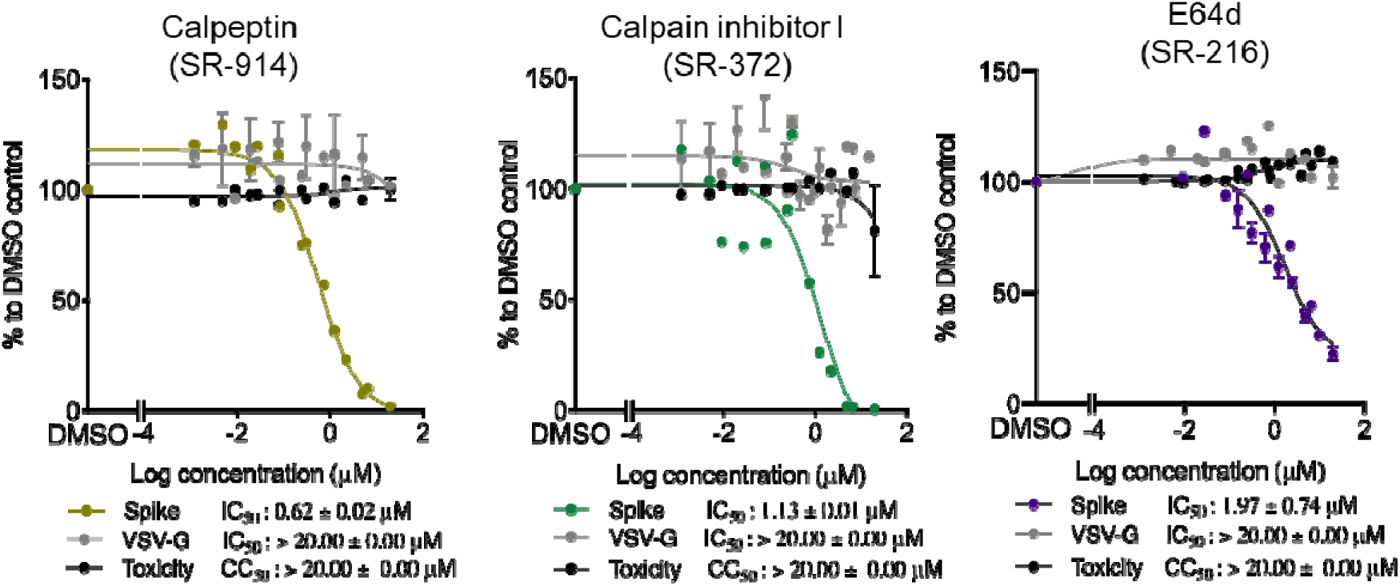
Activity of SR-914, SR-372 and E64d in Vero CCL81 infected with SARS2-S. Cells were incubated with different concentrations of drugs, then infected with SARS2-S or VSV-G. Luciferase was measured 48 hours later, using Bright-Glo. Toxicity was measured, using CellTiter-Glo. E64d was used as control. Shown is the mean ± SEM of n=2 to 5 independent experiments

**Fig.S2.**
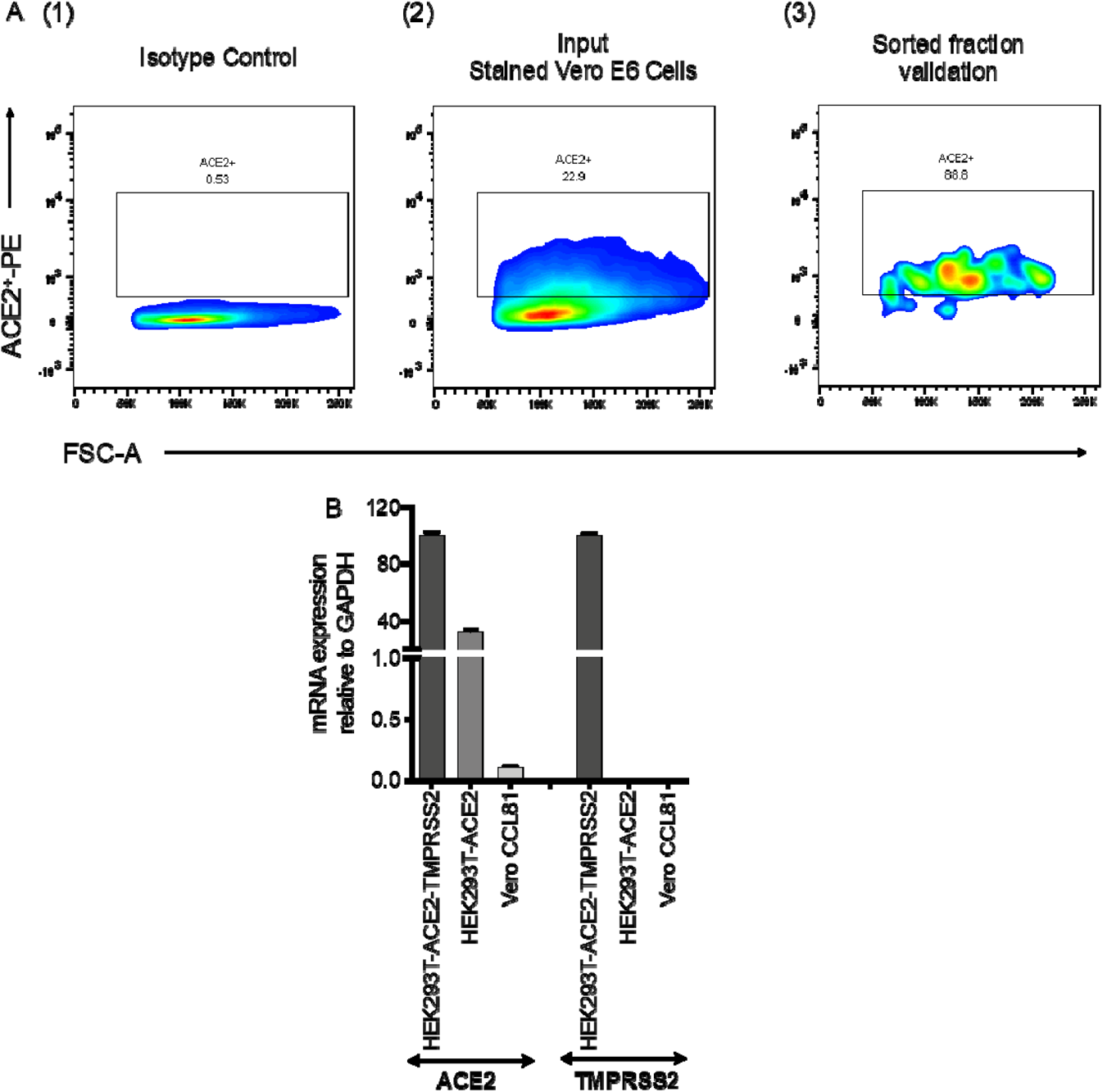
ACE2 and TMPRSS2 expression in different cell lines. **A.** Vero E6 cells were stained with Goat anti-Human Phycoerythrin-conjugated ACE-2 Polyclonal Antibody (R&D Systems, Catalog # FAB933P) for 30 min at 4 °C in dark. Cells were then washed with HBSS and suspended in Pre-Sort buffer (BD Biosciences, Cat#56350). ACE2^+^ cells were gated on the basis of Goat IgG Isotype Control (R&D Systems, Catalog # IC108P) (1), and sorted on FACSAria III. Approximately 23% of the cell population were determined as ACE-positive (2). The cell fraction sorted by the gating was further validated for purity before expansion (3). **B.** Total RNA was extracted, and first-strand cDNA was quantified by qPCR using primers directed to ACE2 or TMPRSS2. Results were normalized as copies of viral mRNA per copy of GAPDH mRNA. The arbitrary value of 100 was assigned to the amount of viral mRNA generated in HEK293T-ACE2-TMPRSS2 cells. Shown is the mean ± SEM of 2 independent experiments.

**Fig.S3.**
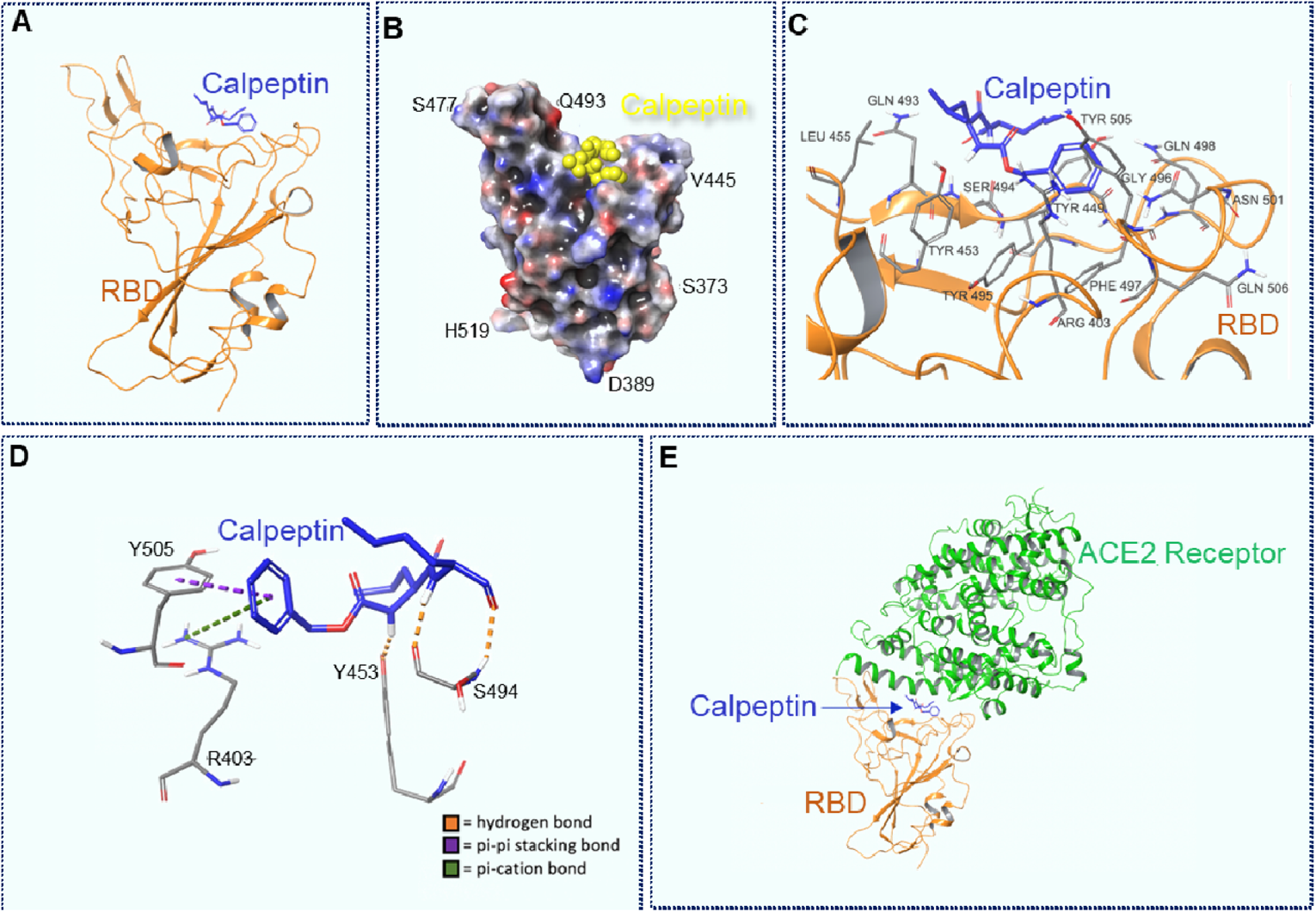
Docking of Calpeptin to the wild type RBD, using a modified PDB 6M0J structure. A. Calpeptin docked to the RBD. B. 3D view of Calpeptin docked to RBD. Molecular surface colored according to residue electrostatic potential. Calpeptin shown in yellow. **C. Calpeptin’s interactions with critical residues within the RBD. D. Types of interaction between Calpeptin and RBD**. Hydrogen bonds are shown in orange, pi-pi stacking bons are shown in purple and pi-cation bonds are shown in green. **E. Calpeptin docked within the RBD-ACE2 connective interface.**

**Fig.S4.**
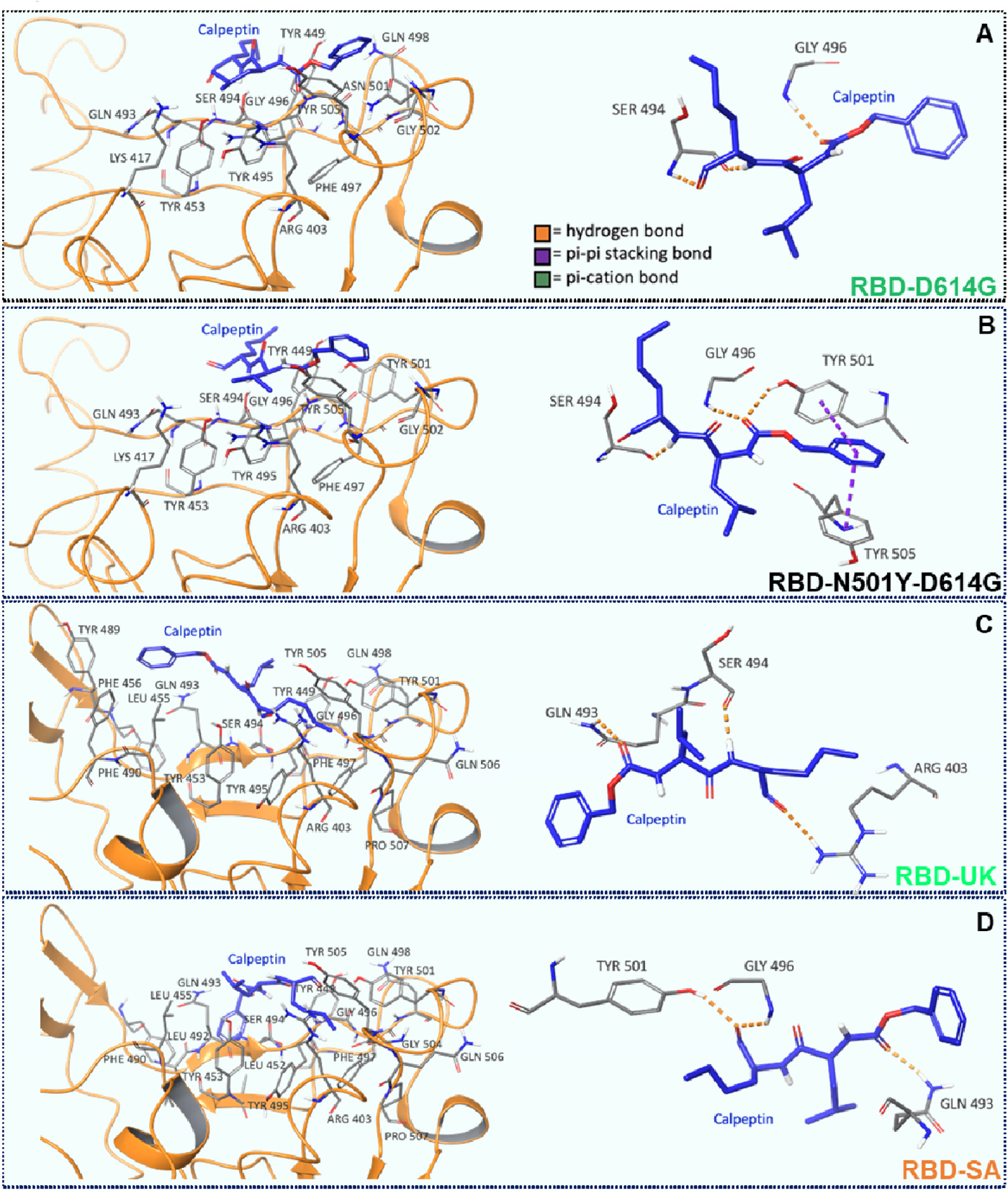
Docking of Calpeptin to the mutants RBD, using a modified PDB 6M0J and 7KDK structures. Calpeptin docked to the D614G, N501Y-D614G, UK and SA variants of SARS-CoV-2 RBD. Residues and types of interactions are shown.

**Fig.S5.**
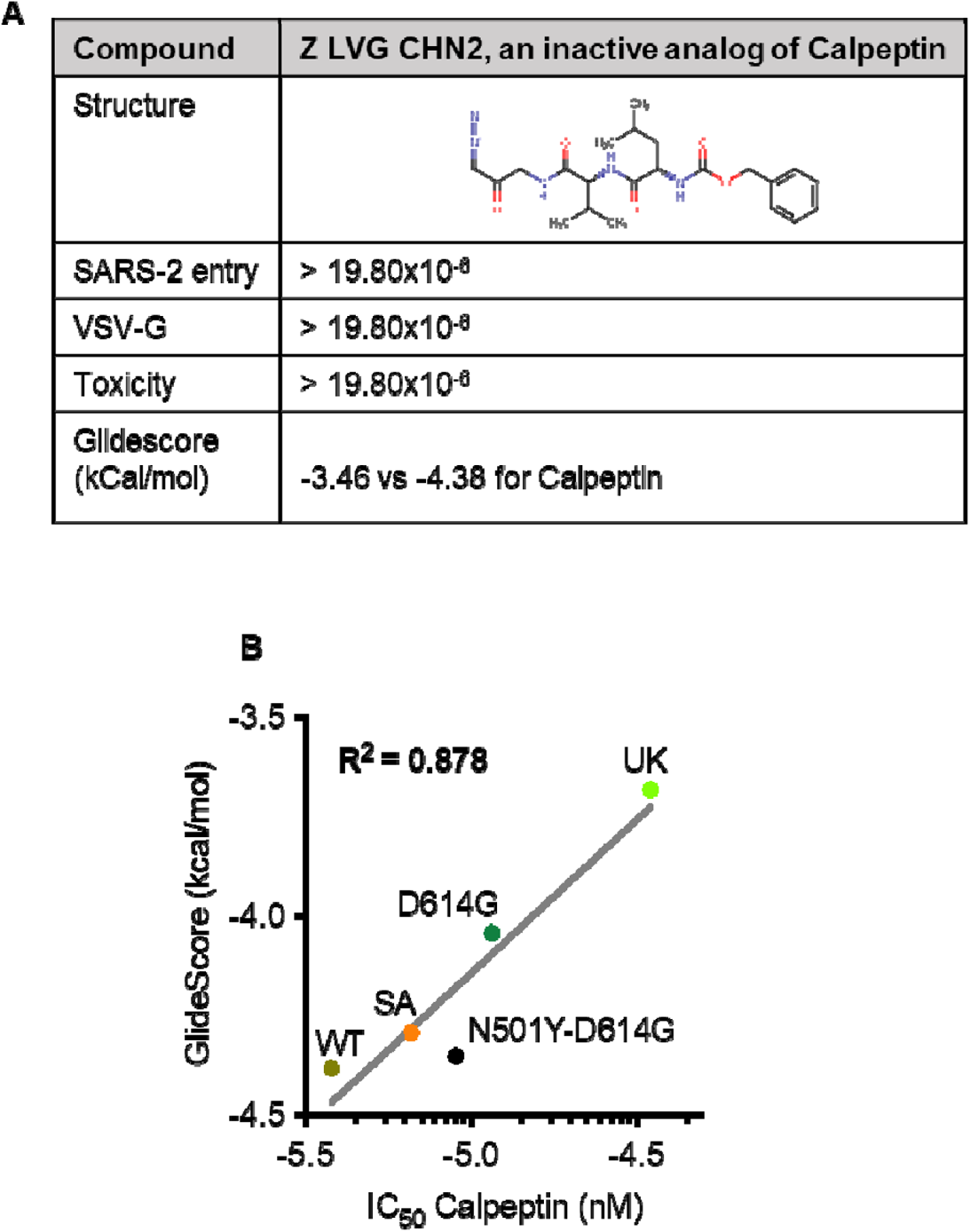
Docking of Calpeptin to RBDs. **A. An inactive analog of Calpeptin binds poorly to wild type RBD**. Z LVG CHN2 was tested in SARS-2 entry assay in HTS. Activity and toxicity were measured. Shown is the mean ± SEM of n=3 independent experiments**. B. Correlation plot between Glidescore and IC50 of Calpeptin against the different mutants of RBD.**

## References

1. Organization, W. H. https://www.who.int/emergencies/diseases/novel-coronavirus-2019. (2020).

2. Mueller, A. L., McNamara, M. S. & Sinclair, D. A. Why does COVID-19 disproportionately affect older people? Aging (Albany NY) 12, 9959–9981, doi:10.18632/aging.103344 (2020).

3. Fu, L. et al. Clinical characteristics of coronavirus disease 2019 (COVID-19) in China: A systematic review and meta-analysis. J Infect 80, 656–665, doi:10.1016/j.jinf.2020.03.041 (2020).

4. Vinayagam, S. & Sattu, K. SARS-CoV-2 and coagulation disorders in different organs. Life Sci 260, 118431, doi:10.1016/j.lfs.2020.118431 (2020).

5. Laskar, P., Yallapu, M. M. & Chauhan, S. C. “Tomorrow Never Dies”: Recent Advances in Diagnosis, Treatment, and Prevention Modalities against Coronavirus (COVID-19) amid Controversies. Diseases 8, doi:10.3390/diseases8030030 (2020).

6. (CDC), T. C. f. D. C. a. P. (2021).

7. Dhama, K. et al. Coronavirus Disease 2019-COVID-19. Clin Microbiol Rev 33, doi:10.1128/CMR.00028-20 (2020).

8. Malin, J. J., Suarez, I., Priesner, V., Fatkenheuer, G. & Rybniker, J. Remdesivir against COVID-19 and Other Viral Diseases. Clin Microbiol Rev 34, doi:10.1128/CMR.00162-20 (2020).

9. Administration, U. S. F. a. D. (2020).

10. Callaway, E. Could new COVID variants undermine vaccines? Labs scramble to find out. Nature 589, 177–178, doi:10.1038/d41586-021-00031-0 (2021).

11. Rubin, R. COVID-19 Vaccines vs Variants—Determining How Much Immunity Is Enough. JAMA, doi:10.1001/jama.2021.3370 (2021).

12. Cherry, J. D. The chronology of the 2002-2003 SARS mini pandemic. Paediatr Respir Rev 5, 262–269, doi:10.1016/j.prrv.2004.07.009 (2004).

13. Wang, Q. et al. Structural and Functional Basis of SARS-CoV-2 Entry by Using Human ACE2. Cell 181, 894–904 e899, doi:10.1016/j.cell.2020.03.045 (2020).

14. Han, Y. & Kral, P. Computational Design of ACE2-Based Peptide Inhibitors of SARS-CoV-2. ACS Nano 14, 5143–5147, doi:10.1021/acsnano.0c02857 (2020).

15. Zamorano Cuervo, N. & Grandvaux, N. ACE2: Evidence of role as entry receptor for SARS-CoV-2 and implications in comorbidities. Elife 9, doi:10.7554/eLife.61390 (2020).

16. Janes, J. et al. The ReFRAME library as a comprehensive drug repurposing library and its application to the treatment of cryptosporidiosis. Proc Natl Acad Sci U S A 115, 10750–10755, doi:10.1073/pnas.1810137115 (2018).

17. Duffy, S. et al. Screening the Medicines for Malaria Venture Pathogen Box across Multiple Pathogens Reclassifies Starting Points for Open-Source Drug Discovery. Antimicrob Agents Chemother 61, doi:10.1128/AAC.00379-17 (2017).

18. Mou, H. et al. Mutations from bat ACE2 orthologs markedly enhance ACE2-Fc neutralization of SARS-CoV-2. bioRxiv, doi:10.1101/2020.06.29.178459 (2020).

19. Moore, M. J. et al. Retroviruses pseudotyped with the severe acute respiratory syndrome coronavirus spike protein efficiently infect cells expressing angiotensin-converting enzyme 2. J Virol 78, 10628–10635, doi:10.1128/JVI.78.19.10628-10635.2004 (2004).

20. Rustanti, L. et al. Differential Effects of Strategies to Improve the Transduction Efficiency of Lentiviral Vector that Conveys an Anti-HIV Protein, Nullbasic, in Human T Cells. Virol Sin 33, 142–152, doi:10.1007/s12250-018-0004-7 (2018).

21. Ramakrishnan, M. A. Determination of 50% endpoint titer using a simple formula. World J Virol 5, 85–86, doi:10.5501/wjv.v5.i2.85 (2016).

22. Baillargeon, P. et al. The Scripps Molecular Screening Center and Translational Research Institute. SLAS Discov 24, 386–397, doi:10.1177/2472555218820809 (2019).

23. Zhang, J. H., Chung, T. D. & Oldenburg, K. R. A Simple Statistical Parameter for Use in Evaluation and Validation of High Throughput Screening Assays. J Biomol Screen 4, 67–73, doi:10.1177/108705719900400206 (1999).

24. Brideau, C., Gunter, B., Pikounis, B. & Liaw, A. Improved statistical methods for hit selection in high-throughput screening. J Biomol Screen 8, 634–647, doi:10.1177/1087057103258285 (2003).

25. Bilinska, K., Jakubowska, P., Von Bartheld, C. S. & Butowt, R. Expression of the SARS-CoV-2 Entry Proteins, ACE2 and TMPRSS2, in Cells of the Olfactory Epithelium: Identification of Cell Types and Trends with Age. ACS Chem Neurosci 11, 1555–1562, doi:10.1021/acschemneuro.0c00210 (2020).

26. Smith, E. et al. High-Throughput Screening for Drugs That Inhibit Papain-Like Protease in SARS-CoV-2. SLAS Discov, 2472555220963667, doi:10.1177/2472555220963667 (2020).

27. Domling, A. & Gao, L. Chemistry and Biology of SARS-CoV-2. Chem 6, 1283–1295, doi:10.1016/j.chempr.2020.04.023 (2020).

28. Riva, L. et al. Discovery of SARS-CoV-2 antiviral drugs through large-scale compound repurposing. Nature 586, 113–119, doi:10.1038/s41586-020-2577-1 (2020).

29. Vitner, E. B. et al. Glucosylceramide synthase inhibitors prevent replication of SARS-CoV-2 and Influenza virus. J Biol Chem 296, 100470, doi:10.1016/j.jbc.2021.100470 (2021).

30. Hoffmann, M. et al. SARS-CoV-2 Cell Entry Depends on ACE2 and TMPRSS2 and Is Blocked by a Clinically Proven Protease Inhibitor. Cell 181, 271–280 e278, doi:10.1016/j.cell.2020.02.052 (2020).

31. Huang, Y., Yang, C., Xu, X. F., Xu, W. & Liu, S. W. Structural and functional properties of SARS-CoV-2 spike protein: potential antivirus drug development for COVID-19. Acta Pharmacol Sin 41, 1141–1149, doi:10.1038/s41401-020-0485-4 (2020).

32. Simmons, G. et al. Inhibitors of cathepsin L prevent severe acute respiratory syndrome coronavirus entry. Proc Natl Acad Sci U S A 102, 11876–11881, doi:10.1073/pnas.0505577102 (2005).

33. Zhou, Y. et al. Protease inhibitors targeting coronavirus and filovirus entry. Antiviral Res 116, 76–84, doi:10.1016/j.antiviral.2015.01.011 (2015).

34. Bosch, B. J., Bartelink, W. & Rottier, P. J. Cathepsin L functionally cleaves the severe acute respiratory syndrome coronavirus class I fusion protein upstream of rather than adjacent to the fusion peptide. J Virol 82, 8887–8890, doi:10.1128/JVI.00415-08 (2008).

35. Drews, K. et al. Glucosylceramide synthase maintains influenza virus entry and infection. PLoS One 15, e0228735, doi:10.1371/journal.pone.0228735 (2020).

36. Baglivo, M. et al. Natural small molecules as inhibitors of coronavirus lipid-dependent attachment to host cells: a possible strategy for reducing SARS-COV-2 infectivity? Acta Biomed 91, 161–164, doi:10.23750/abm.v91i1.9402 (2020).

37. Millet, J. K. & Whittaker, G. R. Physiological and molecular triggers for SARS-CoV membrane fusion and entry into host cells. Virology 517, 3–8, doi:10.1016/j.virol.2017.12.015 (2018).

38. Straus, M. R., Bidon, M., Tang, T., Whittaker, G. R. & Daniel, S. FDA approved calcium channel blockers inhibit SARS-CoV-2 infectivity in epithelial lung cells. bioRxiv, 2020.2007.2021.214577, doi:10.1101/2020.07.21.214577 (2020).

39. Liu, M. et al. Spike protein of SARS-CoV stimulates cyclooxygenase-2 expression via both calcium-dependent and calcium-independent protein kinase C pathways. Faseb j 21, 1586–1596, doi:10.1096/fj.06-6589com (2007).

40. Verrall, G. M. Scientific Rationale for a Bottom-Up Approach to Target the Host Response in Order to Try and Reduce the Numbers Presenting With Adult Respiratory Distress Syndrome Associated With COVID-19. Is There a Role for Statins and COX-2 Inhibitors in the Prevention and Early Treatment of the Disease? Front Immunol 11, 2167, doi:10.3389/fimmu.2020.02167 (2020).

41. Thomas, L. Lipid storm in severe COVID-19 linked to high COX/LOX pathway activit. NEWS MEDICAL LIFE SCIENCE (2020).

42. Chen, J. S. et al. Non-steroidal anti-inflammatory drugs dampen the cytokine and antibody response to SARS-CoV-2 infection. J Virol, doi:10.1128/jvi.00014-21 (2021).

43. Rebecca, V. W. et al. PPT1 Promotes Tumor Growth and Is the Molecular Target of Chloroquine Derivatives in Cancer. Cancer Discov 9, 220–229, doi:10.1158/2159-8290.CD-18-0706 (2019).

44. Wang, M. et al. Remdesivir and chloroquine effectively inhibit the recently emerged novel coronavirus (2019-nCoV) in vitro. Cell Res 30, 269–271, doi:10.1038/s41422-020-0282-0 (2020).

45. Ogando, N. S. et al. SARS-coronavirus-2 replication in Vero E6 cells: replication kinetics, rapid adaptation and cytopathology. J Gen Virol 101, 925–940, doi:10.1099/jgv.0.001453 (2020).

46. Shang, J. et al. Cell entry mechanisms of SARS-CoV-2. Proc Natl Acad Sci U S A 117, 11727–11734, doi:10.1073/pnas.2003138117 (2020).

47. Hoffmann, M. et al. SARS-CoV-2 Cell Entry Depends on ACE2 and TMPRSS2 and Is Blocked by a Clinically Proven Protease Inhibitor. Cell 181, 271–280.e278, doi:10.1016/j.cell.2020.02.052 (2020).

48. Sungnak, W. et al. SARS-CoV-2 entry factors are highly expressed in nasal epithelial cells together with innate immune genes. Nat Med 26, 681–687, doi:10.1038/s41591-020-0868-6 (2020).

49. Riva, L. et al. Discovery of SARS-CoV-2 antiviral drugs through large-scale compound repurposing. Nature 586, 113–119, doi:10.1038/s41586-020-2577-1 (2020).

50. Conn, P. M., Smith, E., Hodder, P., Janovick, J. A. & Smithson, D. C. High-throughput screen for pharmacoperones of the vasopressin type 2 receptor. J Biomol Screen 18, 930–937, doi:10.1177/1087057113483559 (2013).

51. Wan, Y., Shang, J., Graham, R., Baric, R. S. & Li, F. Receptor Recognition by the Novel Coronavirus from Wuhan: an Analysis Based on Decade-Long Structural Studies of SARS Coronavirus. J Virol 94, doi:10.1128/JVI.00127-20 (2020).

52. Gómez, C. E., Perdiguero, B. & Esteban, M. Emerging SARS-CoV-2 Variants and Impact in Global Vaccination Programs against SARS-CoV-2/COVID-19. Vaccines (Basel) 9, doi:10.3390/vaccines9030243 (2021).

53. Lan, J. et al. Structure of the SARS-CoV-2 spike receptor-binding domain bound to the ACE2 receptor. Nature 581, 215–220, doi:10.1038/s41586-020-2180-5 (2020).

54. Zhang, L. et al. SARS-CoV-2 spike-protein D614G mutation increases virion spike density and infectivity. Nat Commun 11, 6013, doi:10.1038/s41467-020-19808-4 (2020).

55. Salleh, M. Z., Derrick, J. P. & Deris, Z. Z. Structural Evaluation of the Spike Glycoprotein Variants on SARS-CoV-2 Transmission and Immune Evasion. Int J Mol Sci 22, doi:10.3390/ijms22147425 (2021).

56. Kumar, S. M. S. Evaluation of the Effect of D614G, N501Y and S477N Mutation in SARS-CoV-2 through Computational Approach. Preprints, doi:10.20944/preprints202012.0710.v1 (2020).

57. Ma, C. et al. Boceprevir, GC-376, and calpain inhibitors II, XII inhibit SARS-CoV-2 viral replication by targeting the viral main protease. Cell Res 30, 678–692, doi:10.1038/s41422-020-0356-z (2020).

58. Siklos, M., BenAissa, M. & Thatcher, G. R. Cysteine proteases as therapeutic targets: does selectivity matter? A systematic review of calpain and cathepsin inhibitors. Acta Pharm Sin B 5, 506–519, doi:10.1016/j.apsb.2015.08.001 (2015).

59. Hopkins, P. L. J. S. Covid-19 Vaccine Makers Take Aim at Dangerous New Strains. The Wall Street Journal (2021).

60. Board, A. (Advisory Board, Advisor Board, Daily Briefing, 2021).

61. Tsujinaka, T. et al. Synthesis of a new cell penetrating calpain inhibitor (calpeptin). Biochem Biophys Res Commun 153, 1201–1208, doi:10.1016/s0006-291x(88)81355-x (1988).

62. Catalano, J. G. et al. Design of small molecule ketoamide-based inhibitors of cathepsin K. Bioorg Med Chem Lett 14, 719–722, doi:10.1016/j.bmcl.2003.11.029 (2004).

63. Catalano, M. & O’Driscoll, L. Inhibiting extracellular vesicles formation and release: a review of EV inhibitors. J Extracell Vesicles 9, 1703244, doi:10.1080/20013078.2019.1703244 (2020).

64. Schneider, M. et al. Severe acute respiratory syndrome coronavirus replication is severely impaired by MG132 due to proteasome-independent inhibition of M-calpain. Journal of virology 86, 10112–10122, doi:10.1128/JVI.01001-12 (2012).

65. Shi, L. et al. [The effect of calpeptin on injury and atrophy of diaphragm under mechanical ventilation in rats]. Zhonghua Wei Zhong Bing Ji Jiu Yi Xue 26, 549–553, doi:10.3760/cma.j.issn.2095-4352.2014.08.005 (2014).

66. Zuo, J. et al. Calpeptin attenuates cigarette smoke-induced pulmonary inflammation via suppressing calpain/IkappaBalpha signaling in mice and BEAS-2B cells. Pathol Res Pract 214, 1199–1209, doi:10.1016/j.prp.2018.06.019 (2018).

67. Tabata, C., Tabata, R. & Nakano, T. The calpain inhibitor calpeptin prevents bleomycin-induced pulmonary fibrosis in mice. Clin Exp Immunol 162, 560–567, doi:10.1111/j.1365-2249.2010.04257.x (2010).

68. Rao, S. S. et al. Calpain-activated mTORC2/Akt pathway mediates airway smooth muscle remodelling in asthma. Clin Exp Allergy 47, 176–189, doi:10.1111/cea.12805 (2017).

69. Liu, Y. et al. Calpain inhibition attenuates bleomycin-induced pulmonary fibrosis via switching the development of epithelial-mesenchymal transition. Naunyn Schmiedebergs Arch Pharmacol 391, 695–704, doi:10.1007/s00210-018-1499-z (2018).

70. Kim, I. U. et al. Antitumor Effect of Calcium-Mediated Destabilization of Epithelial Growth Factor Receptor on Non-Small Cell Lung Carcinoma. Int J Mol Sci 19, doi:10.3390/ijms19041158 (2018).

71. Wang, C. et al. Inhibition of Calpains Protects Mn-Induced Neurotransmitter release disorders in Synaptosomes from Mice: Involvement of SNARE Complex and Synaptic Vesicle Fusion. Sci Rep 7, 3701, doi:10.1038/s41598-017-04017-9 (2017).

72. de la Fuente, S., Sansa, A., Periyakaruppiah, A., Garcera, A. & Soler, R. M. Calpain Inhibition Increases SMN Protein in Spinal Cord Motoneurons and Ameliorates the Spinal Muscular Atrophy Phenotype in Mice. Mol Neurobiol 56, 4414–4427, doi:10.1007/s12035-018-1379-z (2019).

73. Wei, X. et al. Neuroprotective Effect of Calpeptin on Acrylamide-Induced Neuropathy in Rats. Neurochem Res 40, 2325–2332, doi:10.1007/s11064-015-1722-y (2015).

74. Peng, S., Kuang, Z., Zhang, Y., Xu, H. & Cheng, Q. The protective effects and potential mechanism of Calpain inhibitor Calpeptin against focal cerebral ischemia-reperfusion injury in rats. Mol Biol Rep 38, 905–912, doi:10.1007/s11033-010-0183-2 (2011).

75. Mani, S. K. et al. Calpain inhibition preserves myocardial structure and function following myocardial infarction. Am J Physiol Heart Circ Physiol 297, H1744–1751, doi:10.1152/ajpheart.00338.2009 (2009).

76. Froestl, W., Muhs, A. & Pfeifer, A. Cognitive enhancers (nootropics). Part 2: drugs interacting with enzymes. Update 2014. J Alzheimers Dis 42, 1–68, doi:10.3233/jad-140402 (2014).

77. Zanetta, C. et al. Molecular therapeutic strategies for spinal muscular atrophies: current and future clinical trials. Clin Ther 36, 128–140, doi:10.1016/j.clinthera.2013.11.006 (2014).

78. Bordet, T. et al. Identification and characterization of cholest-4-en-3-one, oxime (TRO19622), a novel drug candidate for amyotrophic lateral sclerosis. J Pharmacol Exp Ther 322, 709–720, doi:10.1124/jpet.107.123000 (2007).

79. Iascone, D. M., Henderson, C. E. & Lee, J. C. Spinal muscular atrophy: from tissue specificity to therapeutic strategies. F1000Prime Rep 7, 04, doi:10.12703/p7-04 (2015).

